# Selection leads to false inferences of introgression using popular methods

**DOI:** 10.1101/2023.10.27.564394

**Authors:** Megan L. Smith, Matthew W. Hahn

**Author notes:** **Correspondence** (M.L. Smith).

## Abstract

Detecting introgression between closely related populations or species is a fundamental objective in evolutionary biology. Existing methods for detecting migration and inferring migration rates from population genetic data often assume a neutral model of evolution. Growing evidence of the pervasive impact of selection on large portions of the genome across diverse taxa suggests that this assumption is unrealistic in most empirical systems. Further, ignoring selection has previously been shown to negatively impact demographic inferences (e.g., of population size histories). However, the impacts of biologically realistic selection on inferences of migration remain poorly explored. Here, we simulate data under models of background selection, selective sweeps, balancing selection, and adaptive introgression. We show that ignoring selection sometimes leads to false inferences of migration in popularly used methods that rely on the site frequency spectrum (SFS). Specifically, balancing selection and some models of background selection result in the rejection of isolation-only models in favor of isolation-with-migration models and lead to elevated estimates of migration rates. BPP, a method that analyzes sequence data directly, showed false positives for all conditions at recent divergence times, but balancing selection also led to false positives at medium divergence times. Our results suggest that such methods may be unreliable in some empirical systems, such that new methods that are robust to selection need to be developed.

**Article Summary:** Detecting migration between closely related populations is a central objective in many evolutionary biology studies. However, popular methods for detecting migration assume a simplified model of evolution. Here, we evaluate the impacts of biologically realistic natural selection, recombination, and mutation on three methods for detecting migration. We find that biological complexity leads to false inferences of migration, suggesting that results should be interpreted with caution and that new methods are needed to make robust inferences of migration across empirical systems.

## Introduction

In recent years, as genomic data have become readily available for many taxa, evidence of introgression has accumulated across the tree of life (Mallet *et al*. 2016). A growing interest in understanding the role of introgression in diversification has led to the development of numerous phylogenetic methods for detecting introgression (reviewed in Hibbins and Hahn 2022), but most of these methods cannot detect introgression between sister taxa—only methods that use population genetic data attempt to do this. While detecting introgression between sister taxa is a difficult task, it is of central interest to many researchers. For example, understanding whether closely related taxa exchanged genes during divergence is central to distinguishing among modes of speciation (Payseur and Rieseberg 2016; Roux *et al*. 2016), with evidence of gene flow between closely related taxa sometimes being interpreted as a possible signal of sympatric speciation. Characterizing gene flow between sister species is also essential for developing null models in scans for selection (Williamson *et al*. 2005; Nielsen *et al*. 2007; Excoffier *et al*. 2009; Luqman *et al*. 2021). Thus, despite the difficulties of the task, many population genetic methods have been developed (and have been widely applied) to detect gene flow between sister taxa.

Introgression should lead to increased allele-sharing between taxa and increased variance in coalescence times compared to incomplete lineage sorting (ILS) alone, and methods to detect introgression between sister taxa rely on these expectations. Summary-statistic methods aim to detect particular regions of the genome that have introgressed based on the expectation that these regions should be more similar between sister taxa than non-introgressed regions (Joly *et al*. 2009; Geneva *et al*. 2015; Rosenzweig *et al*. 2016). Other approaches focus on comparing models with and without migration and/or estimating genome-wide migration rates. For example, many site frequency spectrum (SFS)-based methods estimate migration rates and other parameters by finding the parameters that maximize the composite likelihood of the SFS (e.g., Gutenkunst *et al*. 2009; Tellier *et al*. 2011; Excoffier *et al*. 2013), which can be computed using diffusion approximation (e.g., ∂a∂i; Gutenkunst *et al*. 2009) or simulations (e.g., fastsimcoal2; Excoffier *et al*. 2013). Models with and without migration can then be compared based on estimated likelihoods. While powerful, SFS-based approaches do not take advantage of linkage information, and other approaches exist that directly analyze sequence data rather than relying on the SFS as a summary. For example, BPP estimates divergence times and the intensities of introgression events from sequence data under the multispecies coalescent with introgression (MSci) model a using Bayesian Markov chain Monte Carlo (MCMC) approach (Flouri *et al*. 2020). While these methods are powerful and can be highly accurate on simulated datasets, all assume selection does not affect the patterns observed. Several programs attempt to relax this assumption by allowing for variation in effective population sizes across loci, which should mimic some effects of linked selection (e.g., Tine *et al*. 2014; Roux *et al*. 2016; Rougeux *et al*. 2017); however, the efficacy of such programs has not been tested in a wide variety of settings.

Mounting evidence suggests that selection impacts large portions of the genome (reviewed in Cutter and Payseur 2013; Kern and Hahn 2018). Notably, selection can produce genomic signals that mimic demographic processes. For example, linked selection can produce signals that mimic population growth or contraction and can mislead commonly used methods for inferring population size histories (Ewing and Jensen 2016; Schrider *et al*. 2016; Johri *et al*. 2021). Ignoring selection may also pose a substantial problem for methods aiming to detect migration (Cruickshank and Hahn 2014; Mathew and Jensen 2015; Roux *et al*. 2016; Fraïsse *et al*. 2021). Selection leads to increased heterogeneity in levels of divergence among loci by either decreasing (directional selection) or increasing (balancing selection) levels of polymorphism at some loci. Furthermore, balancing selection may maintain polymorphisms for extended periods of time, leading to shared polymorphisms between otherwise diverged species. Since many methods for detecting introgression rely on these same signals, this can lead to false inferences of migration (e.g., Cruickshank and Hahn 2014; Roux *et al*. 2016). Despite more widespread acknowledgement of the role of selection and the shortcomings of neutral assumptions in recent years, methods for inferring migration rates that ignore selection are still widely used.

Here, we simulate data under several evolutionary models that include selection, including background selection, selective sweeps, balancing selection, and adaptive introgression. We evaluate the impact of selection on inferences of migration rates in ∂a∂i, fastsimcoal2, and BPP, and show that some types of selection lead to high rates of false inferences of migration. Our results highlight the importance of incorporating selection into tests for migration.

## Materials and Methods

### Simulations

We simulated two populations that diverged at a set time in the past, *T*_*D*_, and considered two migration histories: no migration (nomig) and a pulse of migration from population 1 to population 2 looking forward in time (p1_p2). We set all population sizes to 125,000 and considered three divergence times: *T*_*D*_ = 0.25 × 4*N*, 1 × 4*N*, and 4 × 4*N* (low, medium, high). The time since introgression, *T*_*M*_, was drawn from a vector {0.01 × *T*_*D*_, 0.05 × *T*_*D*_, 0.10 × *T*_*D*_, 0.15 × *T*_*D*_, …, 0.9 × *T*_*D*_} and the probability of any lineage migrating, *p*_*M*_, was drawn from a vector {0.05, 0.1, 0.15, …, 0.95}. We set the per site mutation rate, *μ*, to 1e-8, and the per site recombination rate, *r*, to 5e-8, both per generation. To lessen the computational burden of forward-in-time simulations, we scaled all simulations by an order of 100: population sizes were scaled to 1250, and mutation rates, recombination rates, and selection coefficients (*s*; see below) were all scaled to keep values of *Nμ*, *Nr*, and *Ns* constant. Similarly, divergence times and the timing of migration were scaled down by an order of 100. To verify that scaling did not bias our results, we also simulated a small number of datasets scaling only by an order of 10 (see Results). We simulated 10,000 independent 10 kb windows for most conditions (see below for details).

To evaluate the impact of selection on inferences of migration rates, we simulated data under six scenarios in SLiM v4.0.1 (Haller and Messer 2019), overlaying neutral mutations with pyslim v1.0.3 and tskit v0.5.5 (Kelleher et al., 2018; Haller et al., 2019). We considered the following selective scenarios: 1) a neutral model; 2) background selection (BGS); 3) a selective sweep in the ancestor of the two populations; 4) a selective sweep in population 1; 5) balancing selection in both populations and their ancestral population; and 6) adaptive introgression. For all scenarios except adaptive introgression, we considered both migration models described above (nomig and p1_p2). For adaptive introgression, we considered only the p1_p2 model. For all simulations, ancestral neutral variation was added via recapitation in pyslim. Each condition is described in detail below:

1. Neutral model: To simulate data in the absence of selection, we overlay all mutations on recorded tree sequences with pyslim.
2. Background selection (“BGS”): To simulate under a model of BGS, we simulated 75% of mutations as deleterious and 25% as selectively neutral. Deleterious mutations had a dominance value of 0.25, corresponding to partially recessive mutations. Selection coefficients for deleterious mutations were drawn from a gamma distribution with a mean and shape of −0.000133 and 0.35 (pre-scaling; mean of −0.0133 post-scaling), corresponding to estimates from *Drosophila* (Huber *et al*. 2017; Schrider 2020). With the population sizes used in our simulations, this corresponds to a mean 2*Ns* = −33.25. We included a burn-in period in which background selection was acting for 25000 generations (post-scaling) prior to population splitting.
3. Selective sweep in the ancestral population (“sweep ancestor”): When simulating a selective sweep in the ancestor of the two populations, the selection coefficient was drawn from a uniform (0.001, 0.005) distribution pre-scaling (0.1, 0.5, post-scaling) with a dominance of 1. At generation 1, a single selectively advantageous mutation was introduced into the ancestral population at position 5000 (i.e. the middle of the locus). Then, until generation 1000 (post-scaling), we checked whether the mutation had fixed or been lost. If it had been fixed, the two populations split at generation 1000 and the simulation proceeded. If the mutation was lost, we restarted at generation 1 and repeated the procedure until the mutation fixed.
4. Selective sweep in population 1 (“sweep p1”): Immediately after divergence, a selectively advantageous mutation with a selection coefficient and dominance as in the linked ancestor simulation was introduced into population 1 at position 5000. If the mutation was lost before migration between populations began (or before the end of the simulation in the no migration model), we restarted the simulation. The mutation had no fitness effect in population 2.
5. Balancing selection (“balancing”): To simulate under a model of balancing selection, we introduced a mutation into the common ancestor of population 1 and population 2 at the beginning of the simulation. We used a mutation effect callback to specify the fitness of this mutation as 1.5 minus the frequency of the mutation in the corresponding population. Thus, when the mutation is rare in a population, it is highly beneficial, but when it is common, it becomes deleterious. Selection should therefore favor maintaining this mutation at an intermediate frequency. At generation 5000 (post-scaling), the two populations split. If the mutation was lost prior to the end of the simulation, we restarted the simulation.
6. Adaptive introgression (“adaptive int”): For this model, only one migration direction was considered (p1_p2). The selection coefficient and dominance were the same as in the model with a selective sweep in P1, except that the mutation was also advantageous in P2. We did not require that the advantageous mutation actually introgressed in these simulations.

We also evaluated the effects of a more biologically realistic model by including variation in mutation rate, recombination rate, and selection coefficients across genomic segments using the “real BGS-weak CNE” approach described in Schrider (2020). Following Schrider (2020), we used annotation data from the University of California Santa Cruz (UCSC) Table Browser for the *D. melanogaster* genome (release 5 / dm3; Adams *et al*. 2000). We also used the *D. melanogaster* recombination map from Comeron *et al*. (2012). Briefly, each simulated replicate was modelled after a randomly selected genomic region with selection coefficients 10-fold smaller in conserved noncoding elements (CNEs) than in coding regions. We modelled windows based on the *Drosophila* genome. For each 10 kb window, we selected an endpoint (constrained to be a multiple of 10 kb). Windows with >= 75% assembly gaps were not allowed, but otherwise windows were drawn randomly with replacement. The locations of annotated exons and phastCons elements were recorded, and deleterious mutations occurred only at these sites. We used recombination rates drawn from the *Drosophila* recombination map for the selected window. We drew the mutation rate from a uniform (3.445e-9, 3.445e-8) distribution (pre-scaling) for each simulated region. We refer to these simulations as the ‘complex genomic architecture’ condition below.

We simulated 10,000 10-kb regions for the background selection and neutral conditions and 1,500 10-kb regions for each sweep condition and balancing selection. These simulated regions were used to build datasets for downstream analyses. We sampled 20 diploid individuals (40 chromosomes) total, 10 per population. We then constructed a genotype matrix, discarding sites that were constant in our sample. We calculated π within each population, F_ST_, and d_xy_ from tree sequences using functions from tskit. We also generated alignments using the generate_nucleotides and convert_alleles functions in pyslim.

These simulated regions were used to construct test datasets (Table 1) for downstream analyses with fastsimcoal2, ∂a∂i, and BPP. Test datasets for background selection and neutral conditions consisted of the 10,000 regions simulated under the corresponding condition and model. For the sweep conditions (sweep p1, sweep ancestor, and adaptive introgression) and balancing selection, we constructed test datasets by sampling 500 (5%), 1000 (10%), or 1500 (15%) regions simulated under the corresponding condition and the remainder of datasets (9500, 9000, or 8500, respectively) from the corresponding set of simulations under a neutral model. Sampling was conducted independently to generate datasets for analyses with SFS-based methods (fastsimcoal2 and ∂a∂i) and BPP. For SFS-based methods, we constructed 100 replicate site frequency spectra (SFS) for each condition by sampling a single SNP per region in the test dataset (see below for additional detail). For analyses with BPP, for each condition, we generated a 500-bp alignment for each region included in the test dataset (see below for additional detail).

**Table 1:** Simulation conditions considered in this study.

| Genomic Architecture | Model | Condition |
| --- | --- | --- |
| uniform | no migration | Neutral<br>Background selection<br>Sweep P1 (5, 10, 15%)<br>Sweep Ancestor (5, 10, 15%)<br>Balancing (5, 10, 15%) |
|  | p1_p2 | Neutral<br>Background selection<br>Sweep P1 (5, 10, 15%)<br>Sweep Ancestor (5, 10, 15%)<br>Balancing (5, 10, 15%)<br>Adaptive Introgression (5, 10, 15%) |
| complex | no migration | Neutral<br>Background selection<br>Sweep P1 (5, 10, 15%)<br>Sweep Ancestor (5, 10, 15%)<br>Balancing (5, 10, 15%) |
|  | p1_p2 | Neutral<br>Background selection<br>Sweep P1 (5, 10, 15%)<br>Sweep Ancestor (5, 10, 15%)<br>Balancing (5, 10, 15%)<br>Adaptive Introgression (5, 10, 15%) |

### Comparing models and estimating migration rates in ∂a∂i

To estimate migration rates in ∂a∂i, we constructed SFS for all simulated datasets (54 datasets per divergence time; Table 1). We built 100 replicate SFS for each dataset, sampling a single bi-allelic SNP with replacement from each simulated fragment (10,000 per dataset). When constructing the SFS for ∂a∂i, we did not populate the monomorphic cell.

We estimated migration rates, population sizes, and divergence times using the split_mig model in ∂a∂i v.2.3.0 (Gutenkunst *et al*. 2009). This model includes two populations, with sizes *V_1_* and *V_2_* relative to the ancestral population, a divergence time, *τ*, and two migration rates, *M_12_* and *M_21_*. We used starting parameter estimates of 0.1 for *V_1_* and *V_2_*, 0.01 for *M_12_* and *M_21_*, and 0.5 *f*or *τ*. We set the lower and upper bounds for *v_1_* and *v_2_* to 1e-3 and 5, respectively. The lower and upper bounds for the migration rate parameters were set to 0 and 5, respectively. For *τ*, the bounds depended on the divergence time of the simulated dataset being analyzed. The upper and lower bounds were set to 0.005 and 1 for low divergence, 0.02 and 4 for medium divergence, and 0.04 and 16 for high divergence. We perturbed parameters using the perturb_params function in ∂a∂i. Then, we optimized parameters using the BOBYQA algorithm, the default algorithm in ∂a∂i. We performed a maximum of 400 evaluations.

For all datasets without migration, we also estimated parameters by maximizing the likelihood of the same model but with migration rates set to zero. Then, we compared the likelihood of the split_mig model to the likelihood of the model without migration using a likelihood ratio test (LRT). We calculated the test statistic Δ as:

Δ = 2 ∗ (*ln* (*split*_*mig*_) − *ln*(*no*_*mig*_)). We then computed the *p*-value using a χ^2^ distribution with two degrees of freedom, and we rejected the null (no migration) model at a significance level of 0.01.

A common approach for accommodating background selection is to allow for different effective population sizes across genomic regions (Roux *et al*. 2016; Rougeux *et al*. 2017). Rougeux *et al*. (2017) developed an approach to accommodate variation in effective population sizes and migration rates across loci in ∂a∂i. They allowed two categories of loci with different effective population sizes for each category, and they allowed for heterogeneous migration across the genome by considering two categories of loci. We ran a modified version of the scripts from Rougeux *et al*. (2017) on our high divergence datasets with a complex genomic architecture and balancing or background selection. We considered five models: SI (strict isolation), SI2N (strict isolation with variation in effective population sizes across loci), IM (isolation with migration), IM2N (isolation with migration with variation in effective population sizes across loci), and IM2m (isolation with migration with variation in migration rates across loci). As above, we optimized parameters using the BOBYQA algorithm and performed a maximum of 400 evaluations. We also adjusted starting parameters for population sizes and divergence times to mirror those used above, except in the IM2m model, for which we used starting values for migration rates of 0.1 and 0.01 for the two different sets of loci. We increased the upper bound on divergence times from the values used by Rougeux *et al*. (2017) to accommodate the deeper divergences simulated in our study. We compared the five models using Akaike Information Criteria (AIC).

### Comparing models and estimating migration rates in fastsimcoal2

To estimate migration rates in fastsimcoal2, we used the SFS constructed for ∂a∂i, except that we included the number of monomorphic sites. To calculate the number of monomorphic sites, we used the following equation:

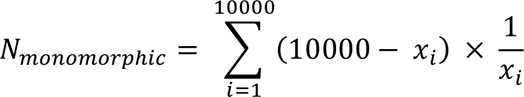

where *x_i_* is the number of segregating sites in fragment *i.* This calculation accounts for the fact that we only sampled a single segregating site per fragment.

We estimated migration rates, population sizes, and divergence times in fastimcoal2 v.2.7.0.9 (Excoffier *et al*. 2013) using a model with the same parameterization as used in ∂a∂i. We set the mutation rate to the value used in the uniform simulations (1e-8). Note that the units used in fastsimcoal2 differ from those used in ∂a∂i. We set the minimum bounds for migration rates to zero, the minimum bounds for population sizes to 1250, and the minimum bounds for the relative population sizes *v_1_* and *v_2_* to 1e-2. The minimum bounds on the divergence time were set to 1250 for low divergence, 5000 for medium divergence, and 10000 for high divergence. We used 100,000 simulations to estimate the expected SFS and performed 40 ECM cycles to estimate parameters. We also compared the migration model to a model with migration rates set to zero using an LRT with two degrees of freedom as described above for ∂a∂i.

### Estimating migration rates with BPP

We estimated introgression probabilities and divergence times in BPP v4.4.0 (Flouri *et al*. 2020). Notably, BPP uses the MSci model, which models an instantaneous introgression event, rather than continuous migration as modelled in our simulations and by ∂a∂i and fastsimcoal2. We first generated sequence alignments for each region equivalent to a 500-bp locus. We used the middle 500 base pairs of each of the 10,000 fragments composing the test dataset. For each simulation condition (Table 1), we created 20 replicate datasets with 500 500-bp loci by sampling loci without replacement from our simulated fragments. In BPP we fixed the species tree to a two-population tree with introgression allowed in both directions and used an inverse gamma (3, 0.01) prior for the parameters θ and τ. Since the inverse gamma prior is a conjugate prior for θ, this allowed the θ parameters to be integrated out analytically, improving run times (Hey and Nielsen 2007). We allowed mutation rates to vary across loci using the a_mubar, b_mubar, and a_mui priors. We set a_mubar and b_mubar to 0, so that mutation rates were relative. We set a_mui equal to 2, and we used the iid prior. We also used the heredity scalar to allow for variation in effective population sizes across loci. For the heredity scalar, we used a Gamma(4,4) prior. We collected 500,000 samples from the posterior after discarding the first 20,000 samples as burnin and sampling every 2 iterations. Some runs did not finish in 90 or 96 hours, but we collected a minimum of 447,060 samples for all runs. These samples were used to estimate all parameters (using the posterior mean). To assess convergence, we used effective sample size (ESS) values. To use the results from BPP to provide a binary determination of the presence of migration, we asked whether the highest posterior density interval (HDI) for migration parameters included 0. An alternative approach is to use Bayes Factors to compare models with and without introgression. The Bayes Factor is the ratio of the marginal likelihoods of two models and requires that we approximate the marginal likelihood of each model. To do this in BPP, we used a path-sampling approach with 8 steps, which required that we run an MCMC algorithm over each step for each model. Given that our MCMC runs take up to 96 CPU hours to complete, this requires ∼1,536 CPU hours per dataset. Because of this, we only calculated BF for a subset of datasets: 5 datasets each from the nomig background selection, nomig neutral, and p1_p2 background selection datasets.

## Results

### Selection alters levels of diversity and divergence

Selection altered patterns of diversity and divergence in both expected and initially surprising ways. Under the model with uniform mutation and recombination rates, the patterns observed in summary statistics conformed to expectations (Supporting Figures S1-S3). Background selection and (to a lesser extent) selective sweeps and adaptive introgression reduced nucleotide diversity relative to the neutral case. Conversely, balancing selection slightly increased diversity. Background selection also decreased divergence between populations, as expected due to reductions in ancestral levels of diversity. Migration from population 1 into population 2 increased nucleotide diversity in population 2 and decreased divergence between populations. Results were more complicated under the model with variation in recombination and mutation rates. As above, selective sweeps and adaptive introgression reduced nucleotide diversity, and balancing selection increased nucleotide diversity relative to the neutral case. However, contrary to our initial expectations, background selection increased nucleotide diversity in many simulations. Such a pattern might be explained by associative overdominance, which maintains neutral diversity when strongly linked to partially recessive deleterious mutations, as in our simulations. Associative overdominance can therefore maintain variation in a population (Ohta 1971; Pamilo and Pálsson 1998; Gilbert *et al*. 2020). To assess whether associative overdominance could be driving our results, we simulated a smaller number of replicates (N=1000) with a dominance coefficient of 0.5, which should eliminate the effects of associative overdominance. As predicted, diversity was reduced relative to the neutral case (Supporting Figure S4). When the scaling of our simulations is less extreme, the amount of nucleotide diversity observed is very similar to the amount observed with more extreme scaling (Figure S4).

We also plotted the SFS under all models and conditions (Supporting Figures S5-S7). As expected, migration led to an increase in the number of variants shared between populations. Balancing selection and background selection with a complex genomic architecture also led to an increase in the number of shared variants between populations, even in the absence of migration. Again, in the background selection case, this could be explained by associative overdominance in regions of low recombination. When the dominance coefficient is set to 0.5, this pattern in the SFS disappears, again supporting associative overdominance as an explanation (Supporting Figure S8). When the scaling of our simulations is less extreme, the SFS results in our background selection simulations remain (Supporting Figure S8D and S8E).

### Selection leads to false inferences of migration using ∂a∂i

Selection sometimes resulted in false inferences of migration in ∂a∂i using a likelihood ratio test (LRT). For the lowest divergence times, false positive rates were elevated even in the absence of selection and were not heavily impacted by selection (Figure 1a,b, Supporting Figure S9). The rates of rejection ranged from 18% to 41%, when only 1% false positives are expected at this *p*-value. When divergence times were moderate, false positive rates were not greater than 1% for any conditions (Figure 1c,d; Supporting Figure S9). Most strikingly, for the highest divergence times considered here, the isolation-only model was often erroneously rejected in favor of the isolation-with-migration model in the presence of balancing and background selection. Under the uniform recombination model, the isolation-only model was rejected in 10, 41, and 50% of replicates with 5, 10, and 15% of loci experiencing balancing selection, respectively (Figure 1e, Supporting Figure S9). Under the complex recombination model, the isolation-only model was rejected in 95, 100, and 100% of replicates with 5, 10, and 15% of loci experiencing balancing selection (Figure 1f, Supporting Figure S9). Background selection led to false positives in 100% of replicates under the complex model (Figure 1f). Under the complex model, selective sweeps also led to false positives in some conditions. Specifically, when 5% of loci experienced sweeps in population 1 or in the ancestor, the isolation-only model was rejected in 28 and 29% of replicates, respectively (Supporting Figure S9). Although the unconstrained model (including migration) should always have a higher likelihood than the constrained model (without migration), this was not always the case. Particularly in the medium divergence case, in ∂a∂i we observed instances in which the constrained model had higher likelihoods, indicating potential issues accurately approximating the likelihood for some datasets (Supporting Figure S10). Perhaps as expected given the LRT results, selection also sometimes resulted in elevated estimates of migration rates in ∂a∂i. As with the LRT, results for the lowest divergence time lack any clear signal related to selection: non-zero rates are observed even in the absence of selection in very recently diverged populations (Figure 2a-b). Again, the most striking impact was seen in the high divergence case in the presence of balancing and background selection (Figure 2e,f, Supporting Figures S11). Migration rates were slightly overestimated under the uniform model with balancing selection. Rates were also overestimated under the complex model with balancing or background selection (Figure 2f), and, in the case of balancing selection, the degree of overestimation increased with the percentage of loci experiencing balancing selection (Supporting Figures S13-S14). When simulations included migration, migration rate estimates tended to be higher under the complex genomic architecture (Supporting Figures S11-S14).

**Figure 1.**
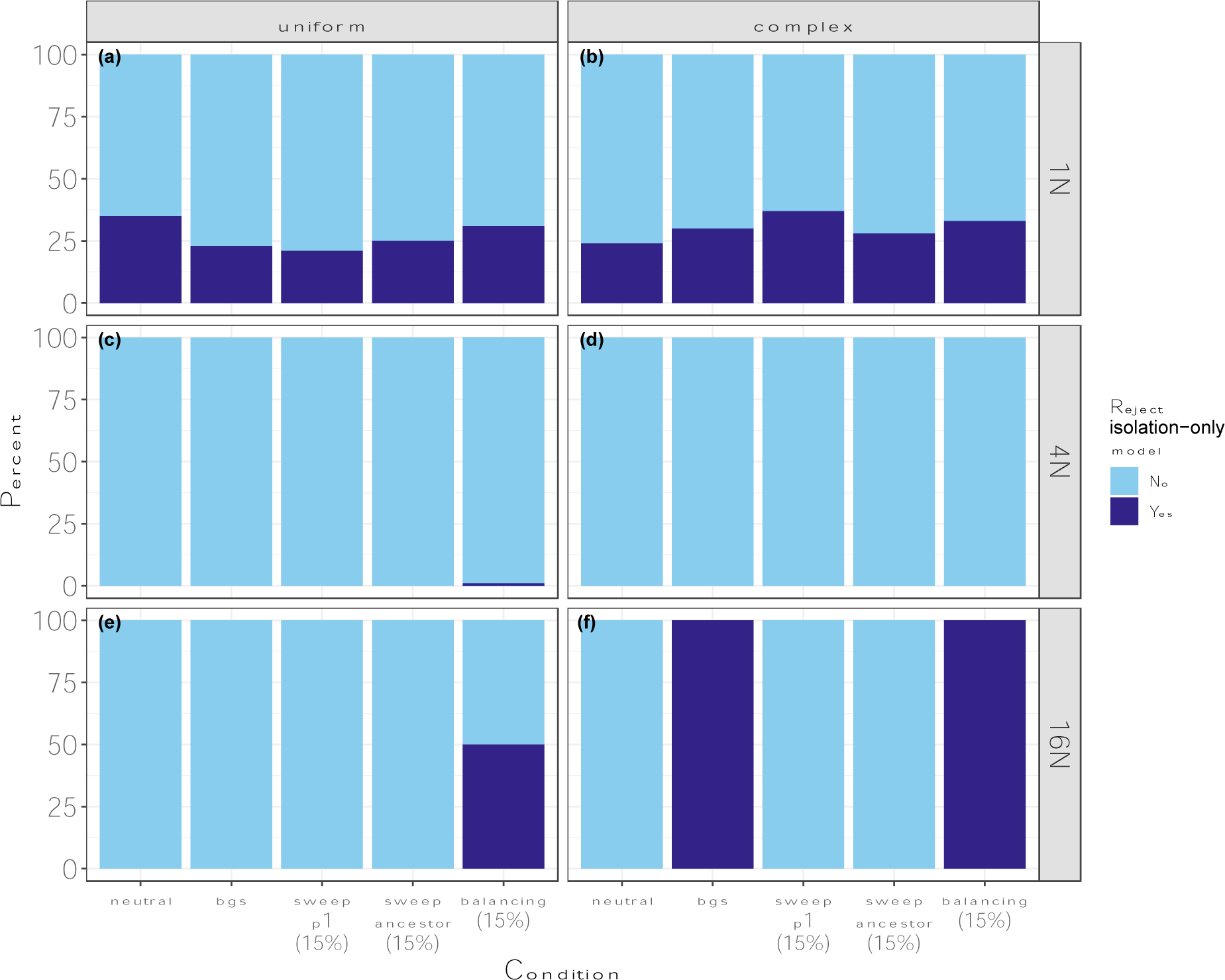
Results of the LRT in ∂a∂i on datasets simulated without migration. Light blue indicates cases where we failed to reject the isolation-only model, and dark blue indicates cases where we rejected the isolation-only model in favor of the isolation-with-migration model. a) results for T=1*N* with a uniform genomic architecture; b) results for T=1*N* with a complex genomic architecture; c) results for T=4*N* with a uniform genomic architecture; d) results for T=4*N* with a complex genomic architecture; e) results for T=16*N* with a uniform genomic architecture; f) results for T=16*N* with a complex genomic architecture.

**Figure 2.**
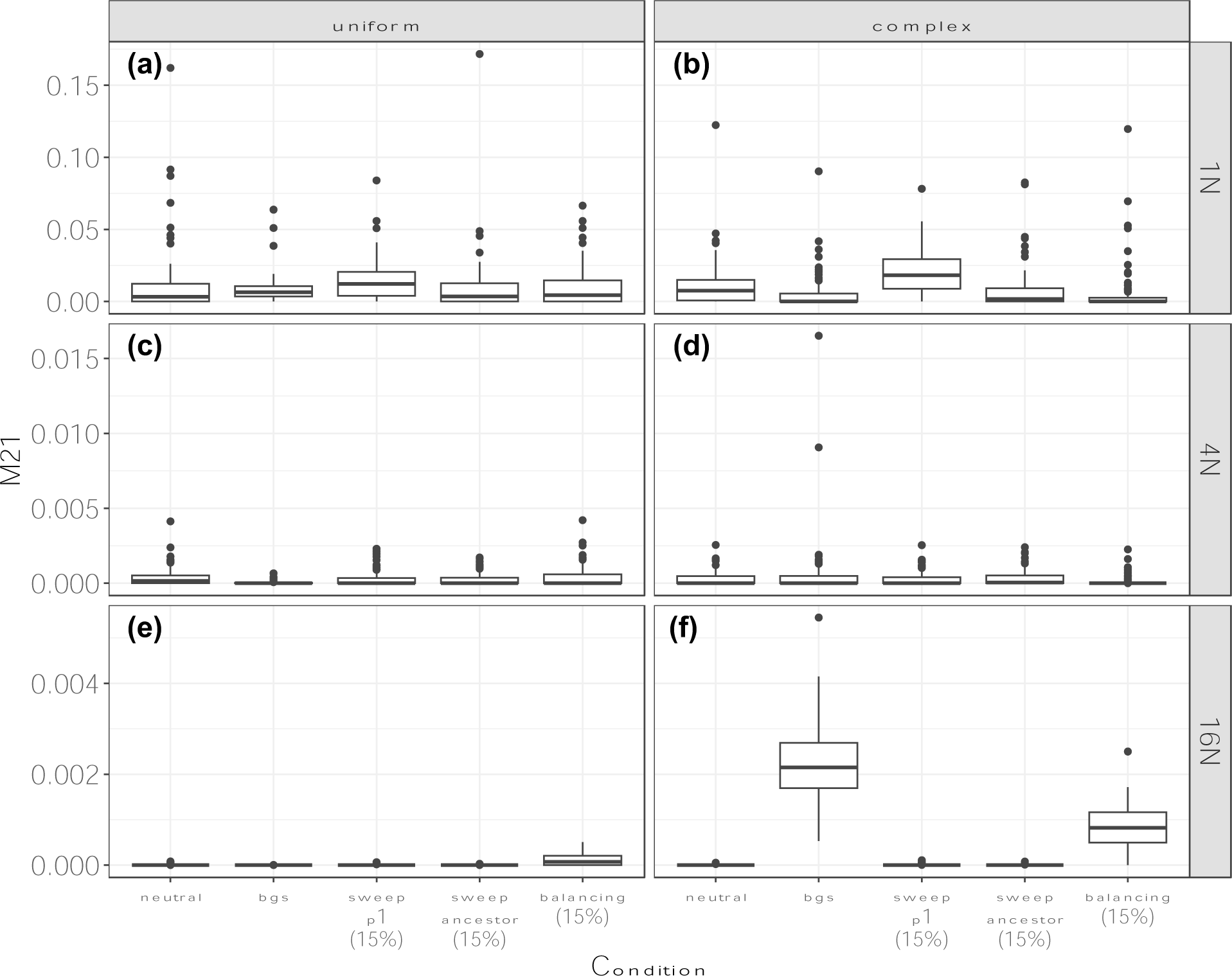
Estimates of *M_21_* in ∂a∂i for datasets simulated without migration. Estimates are in units of 2 × *N*_*ref*_ × *m*_*ij*_. a) results for T=1*N* with a uniform genomic architecture; b) results for T=1*N* with a complex genomic architecture; c) results for T=4*N* with a uniform genomic architecture; d) results for T=4*N* with a complex genomic architecture; e) results for T=16*N* with a uniform genomic architecture; f) results for T=16*N* with a complex genomic architecture.

In the absence of migration and selection, ancestral θ was underestimated (Supporting Figure S15). Background selection under the uniform model and selective sweeps tended to reduce estimates of ancestral θ, while background selection with a complex genomic architecture and balancing selection led to overestimates of ancestral θ when divergence times were high. The presence of migration led to increased estimates of ancestral θ (Supporting Figure S15). We also estimated the size of each population relative to the ancestral population (V_1_, V_2_, Supporting Figures S16-S17). Notably, especially under the uniform genomic architecture, estimates of V_1_ and V_2_ tended to compensate for mistakes in the estimates of ancestral θ. In other words, when ancestral θ was underestimated, V_1_ and V_2_ tended to be overestimated, and vice versa. Divergence time estimates were fairly accurate in the absence of migration and selection, although they were somewhat overestimated in the high divergence case (Supporting Figure S18). When combined with a uniform genomic architecture, background selection and selective sweeps led to overestimates of divergence times in the absence of migration, while balancing selection led to underestimates of divergence times in the high divergence case. However, when combined with a complex genomic architecture and moderate or high divergence times, background selection led to underestimated divergence times. Divergence times were always underestimated in the presence of migration, except for at the lowest divergence times with background selection and a uniform genomic architecture.

For the two cases with the highest false positive rates (background and balancing selection with high divergence times and a complex genomic architecture), we compared models with and without migration, with and without variation in effective population sizes, and with or without variation in migration rates following Rougeux *et al*. (2017). With background selection, this led to a reduction in false positives (from 100% to 81%), but for the majority of replicates, a model with migration was still selected as the best model, and the most commonly selected model included two categories of migration rates (Supporting Figure S19a). For balancing selection, this approach also reduced the false positive rate (from 100% to 83%), and the isolation-with-migration model was selected most often (Supporting Figure S19b).

### Selection leads to false inferences of migration using fastsimcoal2

Background and balancing selection often resulted in false inferences of migration in fastsimcoal2. For the lowest divergence times, we rarely rejected the isolation-only model (Figure 3a-b, Supporting Figure S20). When divergence times were medium, false positive rates were elevated across all conditions, even in the absence of selection: rates of rejection ranged from 13% to 45% (Figure 3c-d). For the highest divergence times considered here, the isolation-only model was always erroneously rejected in favor of the isolation-with-migration model in the presence of background selection and a complex genomic architecture (Figure 3f). Under the complex genomic architecture, the isolation-only model was rejected in 21, 81, and 100% of replicates with balancing selection in 5, 10, and 15% of loci, respectively. (Figure 3f, Supporting Figure S20). There was also a slightly elevated false positive rate (4%) under a uniform genomic architecture when 15% of loci experienced balancing selection (Figure 3e). Although the unconstrained model (including migration) should always have a higher likelihood than the constrained model (without migration), as with ∂a∂i this was not always the case. Particularly in the low divergence case, in fastsimcoal2 we observed instances in which the constrained model had higher likelihoods, indicating potential issues accurately approximating the likelihood (Supporting Figure S21).

**Figure 3.**
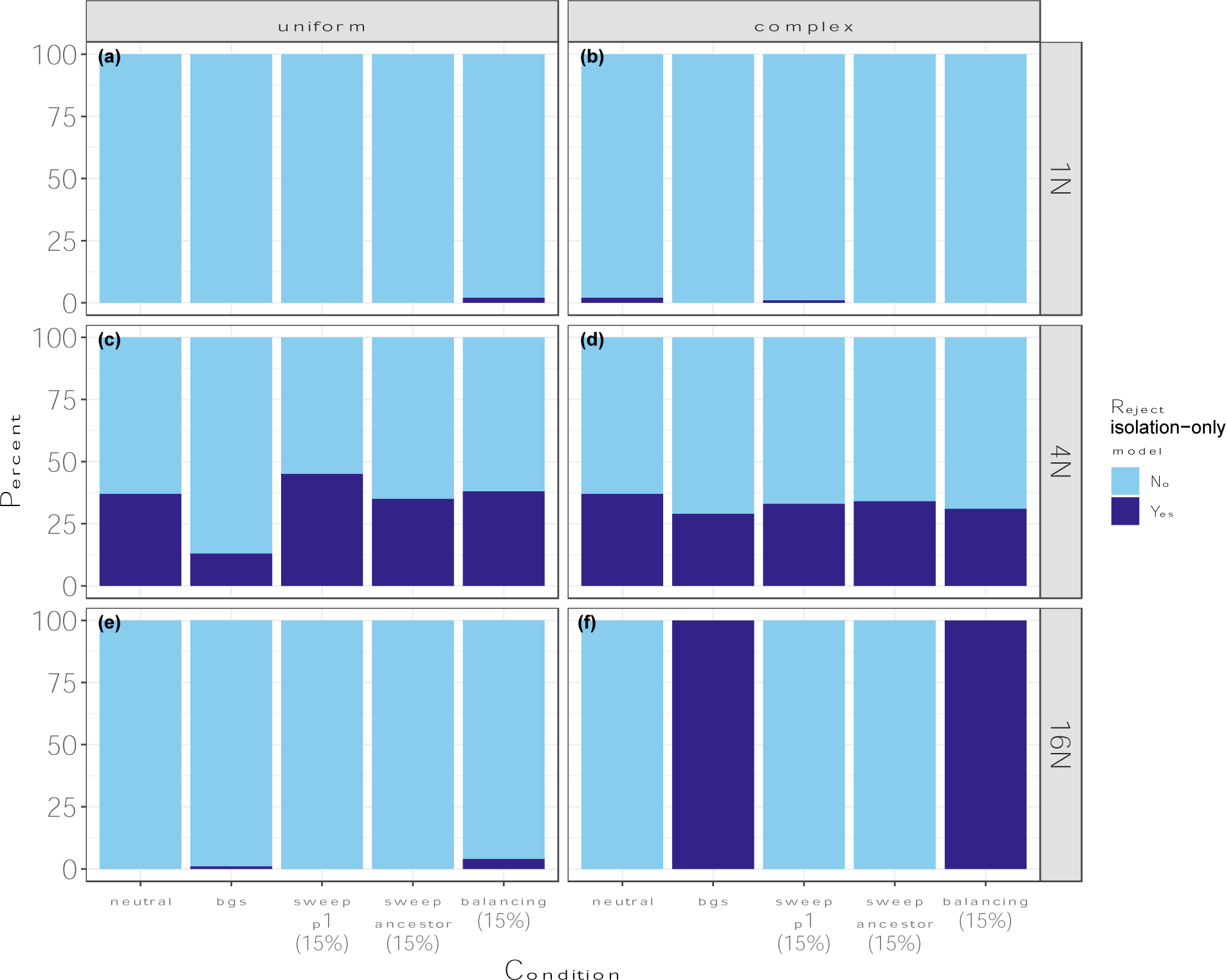
Results of the LRT in fastsimcoal2 for datasets simulated without migration. Light blue indicates cases where we failed to reject the isolation-only model, and dark blue indicates cases where we rejected the isolation-only model in favor of the isolation-with-migration model. a) results for T=1*N* with a uniform genomic architecture; b) results for T=1*N* with a complex genomic architecture; c) results for T=4*N* with a uniform genomic architecture; d) results for T=4*N* with a complex genomic architecture; e) results for T=16*N* with a uniform genomic architecture; f) results for T=16*N* with a complex genomic architecture.

As with ∂a∂i, selection resulted in elevated estimates of migration rates in fastsimcoal2 (Figure 4, Supporting Figures S22-S25). When divergence times were high, migration rates were slightly overestimated with a uniform genetic architecture and balancing selection and were substantially overestimated in the presence of a complex genomic architecture and background or balancing selection (Figure 4f). In the case of balancing selection, the degree of overestimation increased with the percentage of loci experiencing balancing selection (Supporting Figures S24-S25). When simulations included migration, migration rate estimates tended to be lower under the complex genomic architecture and were slightly elevated in the presence of background selection with a complex genomic architecture, balancing selection, and adaptive introgression (Supporting Figures S22-S25).

**Figure 4.**
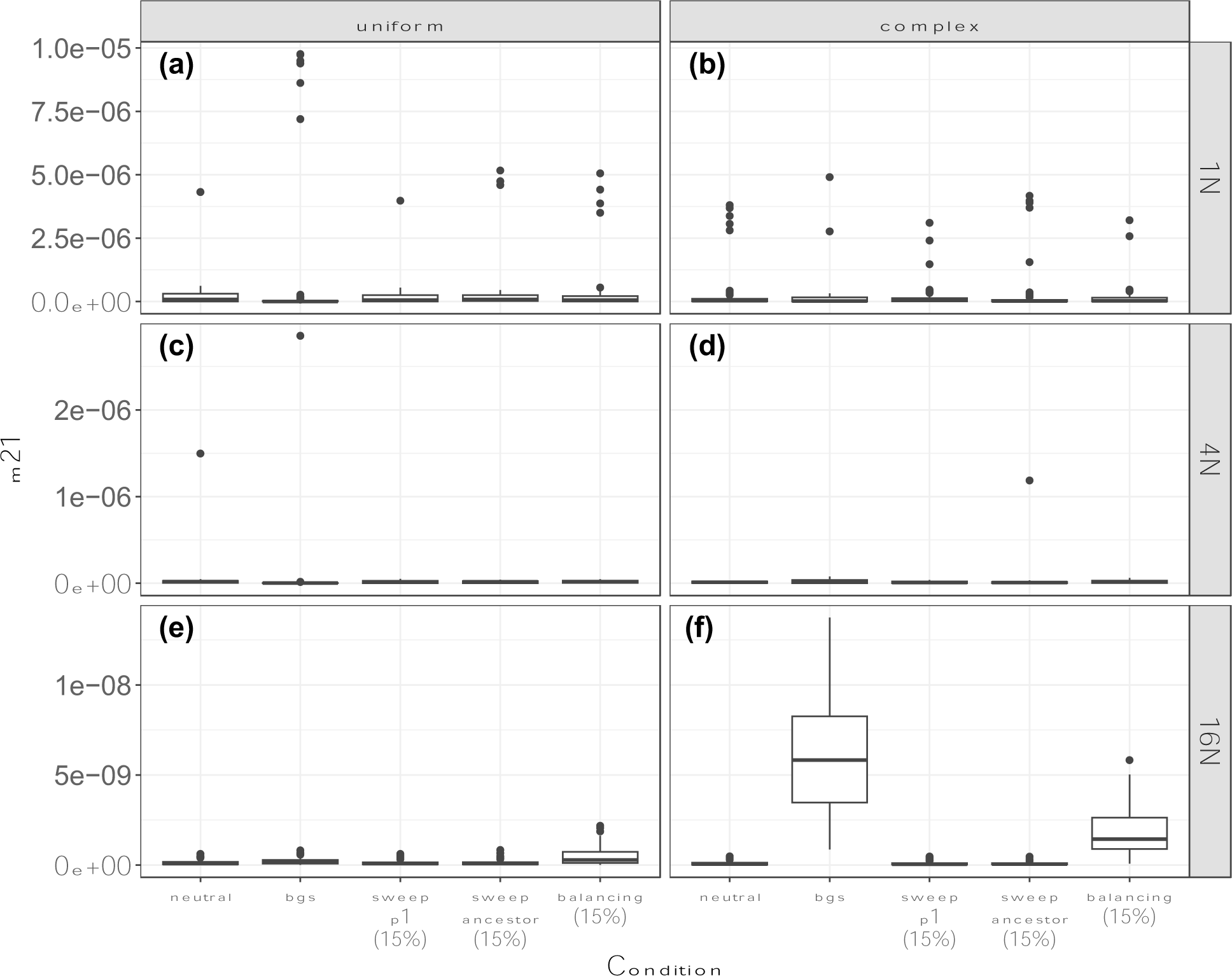
Estimates of *m_21_* in fastsimcoal2 for datasets simulated without migration. The rate *m_ij_* is the probability of any gene moving from population *i* to population *j* backwards in time each generation. a) results for T=1*N* with a uniform genomic architecture; b) results for T=1*N* with a complex genomic architecture; c) results for T=4*N* with a uniform genomic architecture; d) results for T=4*N* with a complex genomic architecture; e) results for T=16*N* with a uniform genomic architecture; f) results for T=16*N* with a complex genomic architecture.

Estimates of the ancestral population sizes were fairly accurate in fastsimcoal2 (Supporting Figure S26). The relative population sizes V_1_ and V_2_ were overestimated across all models and conditions (Supporting Figures S27-S28). In the absence of selection, migration, and variation in mutation and recombination rates divergence time estimates were accurate (Supporting Figure S29). Divergence time estimates were higher under the complex genomic architecture compared to the uniform genomic architecture and were reduced in the presence of background selection (Supporting Figure S29). Divergence times tended to be underestimated in the presence of migration (Supporting Figure S29).

### Selection leads to false inferences of migration in BPP

Using BPP, non-zero migration rates were often inferred in the absence of migration (Figure 5; Supporting Figures S30-S32). The highest posterior density interval (HDI) of the two migration parameters, φ*-X* and φ*-Y*, rarely contained zero in the low divergence case, regardless of the presence of selection (Figure 5a,b; Supporting Figure S30). In the medium and high divergence cases, BPP still inferred non-zero migration often. In the medium divergence case, under the uniform recombination and mutation model, the HDI for φ*-Y* did not include zero in 30, 30, and 40% of replicates with balancing selection in 5, 10, and 15% of loci (Figure 5c; Supporting Figure S31). Similarly, the HDI for φ*-X* did not include zero in 10, 35, and 45% of replicates with balancing selection in 5, 10, and 15% of loci (Figure 5c; Supporting Figure S32). Under the complex model, the HDI for φ*-Y* did not include zero in 10, 25 and 40% of replicates with balancing selection in 5, 10, and 15% of loci (Figure 5d; Supporting Figure S31). Similarly, the HDI for φ*-X* did not include zero in 25, 45, and 65% of replicates (Figure 5d; Supporting Figure S32). False positive rates were also slightly elevated across other conditions in the medium divergence case (0-10% false positive rate). (Figure 5d; Supporting Figures S30-S32). In the high divergence case, BPP inferred non-zero migration in 0-10% of replicates under each model and condition, but there was no clear pattern with respect to selection and genomic architectures (Figure 5e-f; Supporting Figures S30-S32). Although Bayesian methods do not have an equivalent “false positive” rate to frequentist methods, we would not expect such a high proportion of HDIs to not include the true parameter value. We also used Bayes Factors to compare a model with introgression to a model without introgression for a subset of datasets with high divergence (Supporting Table S1). We rejected the model without migration in 40% of replicates without migration under both neutral and background selection conditions, and we failed to reject the model without migration in 20% of replicates with migration and background selection.

**Figure 5.**
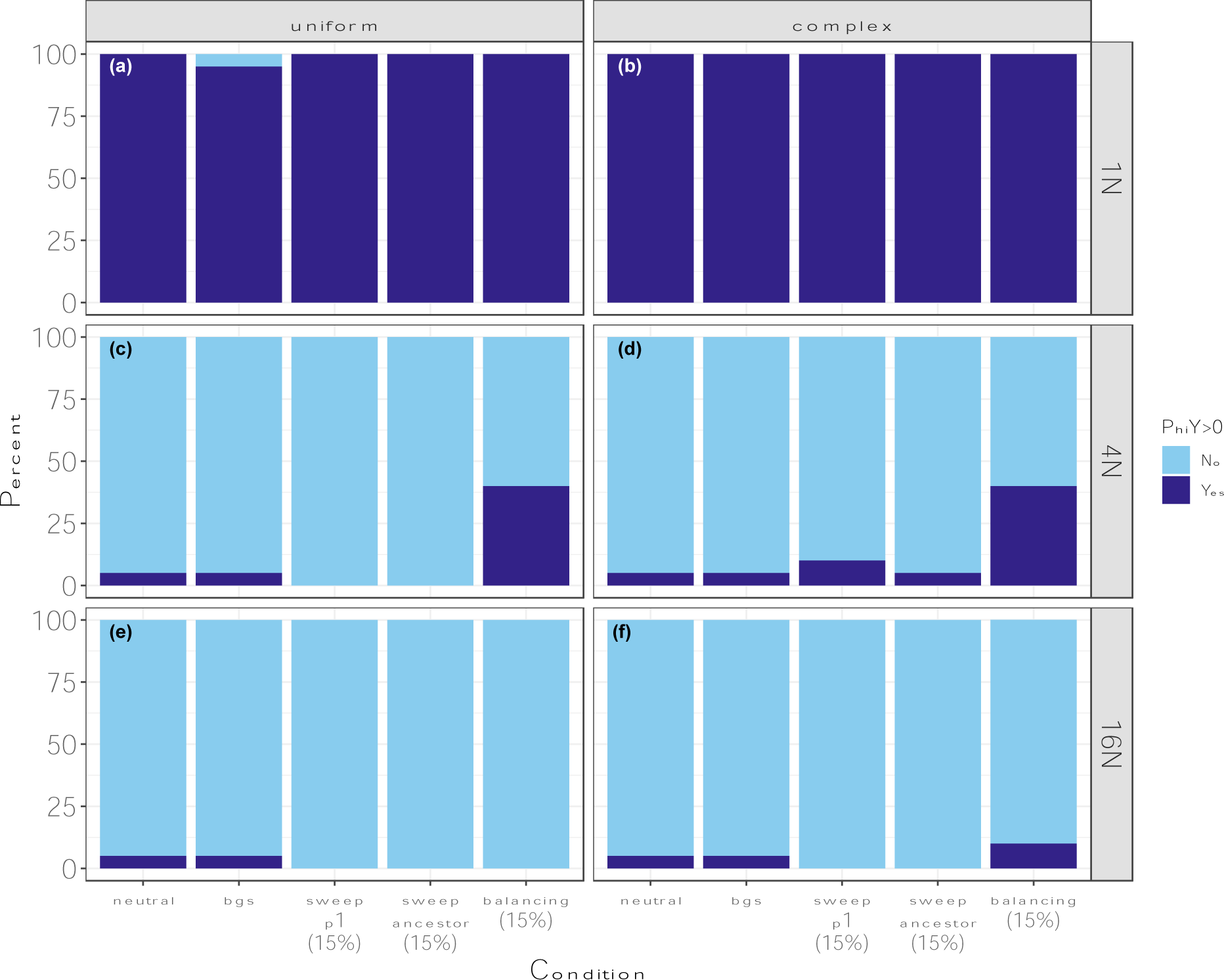
The proportions of replicates with 95% highest posterior density intervals for φ*-Y* that do not include zero in BPP for datasets simulated without migration. Light blue indicates cases where the HDI includes zero, and dark blue indicates cases where the HDI does not include zero. a) results for T=1*N* with a uniform genomic architecture; b) results for T=1*N* with a complex genomic architecture; c) results for T=4*N* with a uniform genomic architecture; d) results for T=4*N* with a complex genomic architecture; e) results for T=16*N* with a uniform genomic architecture; f) results for T=16*N* with a complex genomic architecture.

As expected, given that the HDIs do not overlap zero, migration rate estimates were elevated in the presence of balancing selection (Figure 6; Supporting Figures S33). When simulations included migration, estimates of φ*-X* and φ*-Y* were generally higher under the complex genomic architecture (Supporting Figures S33-S34). Results were qualitatively similar whether 5, 10, or 15% of sweep datasets experienced a sweep (Supporting Figures S35-S36).

**Figure 6.**
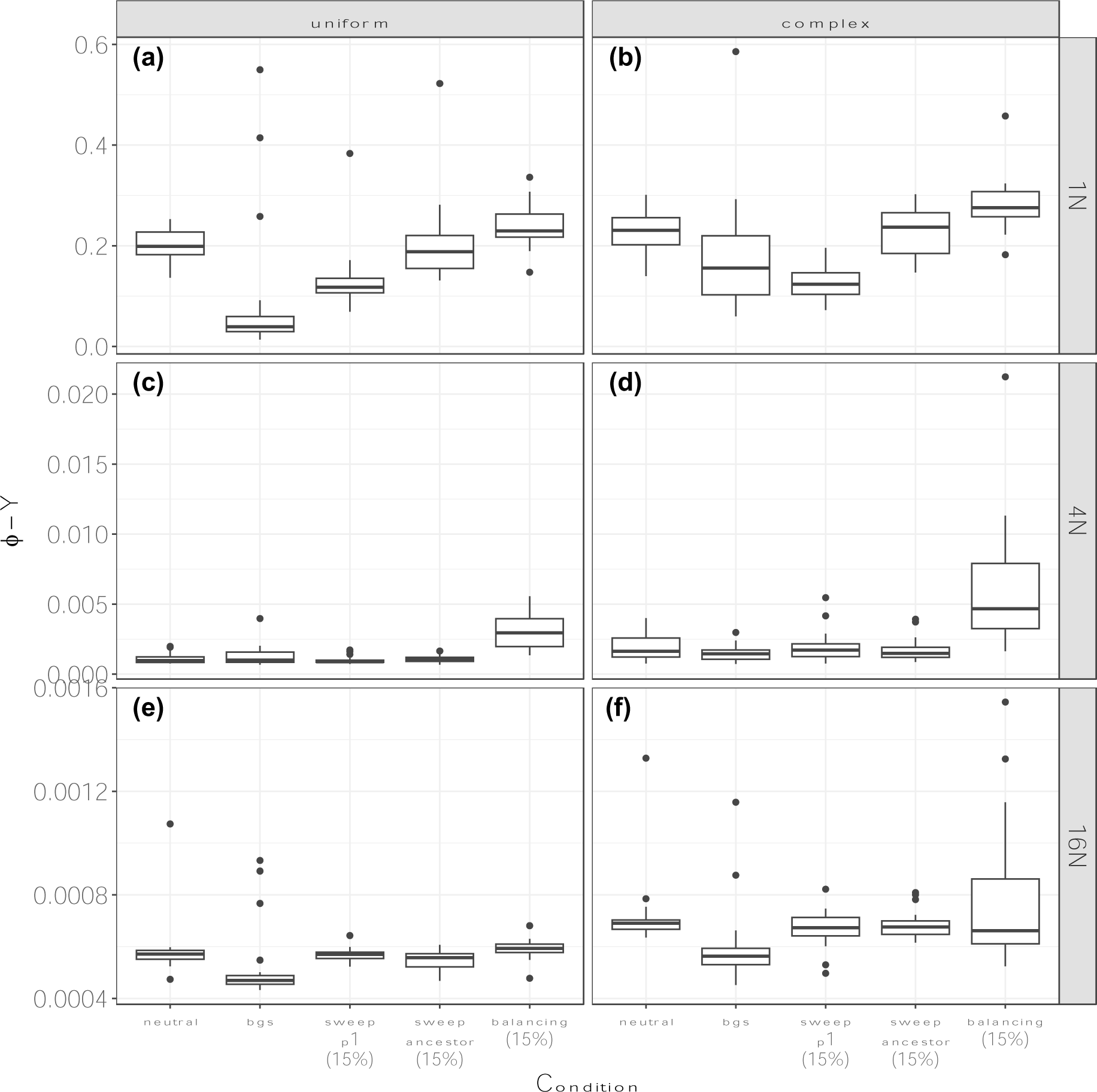
Mean posterior estimates of φ*-Y* in BPP for datasets simulated without migration. φ*-Y* _is_ the weight of the introgression edge Y in the MSci model. a) results for T=1*N* with a uniform genomic architecture; b) results for T=1*N* with a complex genomic architecture; c) results for T=4*N* with a uniform genomic architecture; d) results for T=4*N* with a complex genomic architecture; e) results for T=16*N* with a uniform genomic architecture; f) results for T=16*N* with a complex genomic architecture.

We also evaluated whether there was evidence for a lack of convergence in BPP runs by examining ESS values. We focused on ESS values for the log likelihood, along with the φ*-X* and φ*-Y* parameters. In the low divergence case, ESS values for the φ parameters were often low, indicating a lack of convergence, particularly under the complex genomic architecture; however, ESS values were generally greater than 200 in the medium and high divergence cases (Supporting Figure S37). Under some conditions, there was evidence of a correlation between ESS values and parameter estimates (e.g., for φ*-X* under a neutral model with a complex genomic architecture, Supporting Figure S38). This suggests that in some (but not all) instances, assessing convergence may allow researchers to identify problematic cases.

In the absence of selection and migration, divergence times were generally overestimated (Supporting Figure S39). Divergence time estimates were reduced in the presence of background selection and were elevated under the complex genomic architecture relative to the uniform genomic architecture. The presence of migration led to underestimates of divergence times when divergence times were high.

## Discussion

Our results suggest that, while popular methods for estimating migration rates between sister populations or species are largely robust to selective sweeps and simple models of background selection, models with non-uniform recombination rates and models with balancing selection can lead to high rates of false positives. The three approaches tested (fastsimcoal, ∂a∂i, and BPP) all showed high rates of misleading results in the presence of balancing selection and background selection with a complex genetic architecture (Figures 1, 3, 5). Given that large portions of the genomes of many species are impacted by selection (Begun *et al*. 2007; McVicker *et al*. 2009; Sella *et al*. 2009; Langley *et al*. 2012; Corbett-Detig *et al*. 2015; Phung *et al*. 2016; Pouyet *et al*. 2018) and that variation in mutation and recombination rates across the genome is the norm, these results suggest that some inferences of introgression may be artifacts that do not reflect biological reality.

Numerous studies have found that ignoring natural selection can negatively impact different types of demographic inferences. Selection can lead to false inference of population size changes (e.g., Ewing and Jensen 2016; Schrider *et al*. 2016; Johri *et al*. 2021) and several studies have suggested that selection can also mislead inferences of migration (e.g., Cruickshank and Hahn 2014; Mathew and Jensen 2015; Roux *et al*. 2016). In our study, false inferences appear to be primarily driven by the impacts of balancing selection and associative overdominance. Both balancing selection and associative overdominance increase the number of polymorphisms shared between two populations. This effect is clearly visible in the SFS produced under these conditions, which resemble those produced by migration (Supporting Figures S6-S7). It remains unclear how prevalent these patterns are likely to be in empirical systems. Notably, we did not set out to simulate the effects of associative overdominance—our simulations were parameterized based on the *Drosophila melanogaster* genome, and this unexpectedly led to associative overdominance. In species with compact genomes and low recombination regions, therefore, we may expect these impacts to potentially be widespread. In species with large genomes, but many functional non-coding elements, the impacts may also extend throughout the genome (e.g., Gilbert *et al*. 2020). Furthermore, while the balancing selection simulated here may seem extreme, there are numerous examples of trans-specific polymorphisms maintained by balancing selection (e.g., the *S*-locus in flowering plants (Wright 1939; Le Veve *et al*. 2023)). We recommend that researchers exclude such loci and neighboring regions—if they can be identified—when conducting demographic inference.

To account for the effects of selection, several approaches for inferring migration have been developed that allow for heterogeneous effective population sizes among loci (Sousa *et al*. 2013; Roux *et al*. 2016; Sethuraman *et al*. 2019; Fraïsse *et al*. 2021). Although these methods vary in the types of inferences that can be made—from locus-specific migration rates to genome-wide migration rates—they all model selection by allowing for variation in θ among loci. Notably, while this may accommodate some of the simpler effects of background selection (e.g., reduced diversity and increased variation in coalescence times), it is unlikely to accommodate the impacts of associative overdominance or balancing selection. To evaluate this, we applied such an approach to compare models in ∂a∂i, and, while false positive rates were reduced, they were still high (81% and 83% for background and balancing selection, respectively). BPP also allows for variation across loci, either by using a rate multiplier for θ and τ or for θ alone (Flouri *et al*. 2020). In our analyses, we used both rate multipliers and still found relatively high false positive rates, particularly in the presence of balancing selection. It is not clear that the results using any other similar methods would differ qualitatively from these (to our knowledge, none have been tested against a no-migration scenario with selection).

Phylogenetic methods for inferring gene flow are much more robust to assumptions about selection, largely because they often depend on asymmetries in tree topologies (Hibbins and Hahn 2022). However, the dependence on tree asymmetry also means that they cannot be used to detect gene flow between sister lineages. So, what is the way forward? Our results, along with previous studies, highlight several potential possibilities. Statistical-learning approaches are highly flexible (Schrider and Kern 2018), and one path forward involves training these algorithms under appropriate models that incorporate selection. One such approach has successfully used approximate Bayesian computation to jointly estimate DFEs and population size histories (Johri *et al*. 2020). Beyond training statistical learning algorithms on more realistic training data, techniques for domain adaptation—a subfield of machine learning that aims to adapt an algorithm trained on the source domain (e.g., on simulations under a model of interest) to the target domain (i.e., empirical data; reviewed in Wilson and Cook 2020) offer a promising path forward to accommodating complex biological realities in population genomics (e.g., Mo and Siepel 2023). Regardless of which methods are used, accurate inferences of demographic histories will have to include the complexities introduced by selection.

Accurate inferences of introgression histories are important because they can tell us about modes of speciation. A large number of studies supporting gene flow between closely related populations have been interpreted as lending support to speciation-with-gene-flow models; our results highlight that caution is warranted in these interpretations (cf. Cruickshank and Hahn, 2014), though there are other reasons to exercise caution as well (Yang *et al*. 2017). Although many estimates were non-zero, migration rates estimated for datasets generated under models including selection were low absolutely. In fastsimcoal2, estimated migration rates per generation were on the order of 10^-9^, in ∂a∂i, 2 × *N*_*ref*_ × *m*_*ij*_ was on the order of 0.002 (∼10^-9^ migrants per generation), and in BPP the weight of the hybrid edge was on the order of 0.002 (Figures 2, 4, 6). On the other hand, estimates of effective population size often change multiple orders of magnitude over extremely short periods of time in analyses of empirical data using these same methods (e.g., Rosser *et al*. 2024), suggesting that there are other biological complexities not captured by these methods (or by our simulations). Moving forward, we recommend that inferences of migration between sister populations made without considering selection be interpreted with caution, particularly when inferred rates of migration are low. Importantly, we do not believe that the results found here with a limited set of selective scenarios and a limited set of introgression histories can fully describe the effects of selection, mutation, and recombination on inaccurate demographic inferences. Inferences about the presence, direction, and timing of introgression (including whether speciation and introgression occur at the same time—i.e. homoploid hybrid speciation) may all be affected by models that ignore natural selection and complex genomic architectures. We hope that new methods can be developed to overcome these obstacles.

## Data Availability

Simulated data formatted for various programs are available on Figshare (DOI: 10.6084/m9.figshare.24354277). All scripts are available on GitHub (https://meganlsmith.github.io/selectionandmigration/).

## Acknowledgements

We thank Peter Ralph and two reviewers for very helpful comments and suggestions. This work was supported by a National Science Foundation (NSF) postdoctoral fellowship to MLS (DBI-2009989) and NSF grant to MWH (DBI-2146866).

