## Supporting Information for "Selection leads to false inferences of introgression using popular methods"

### Supporting Figures

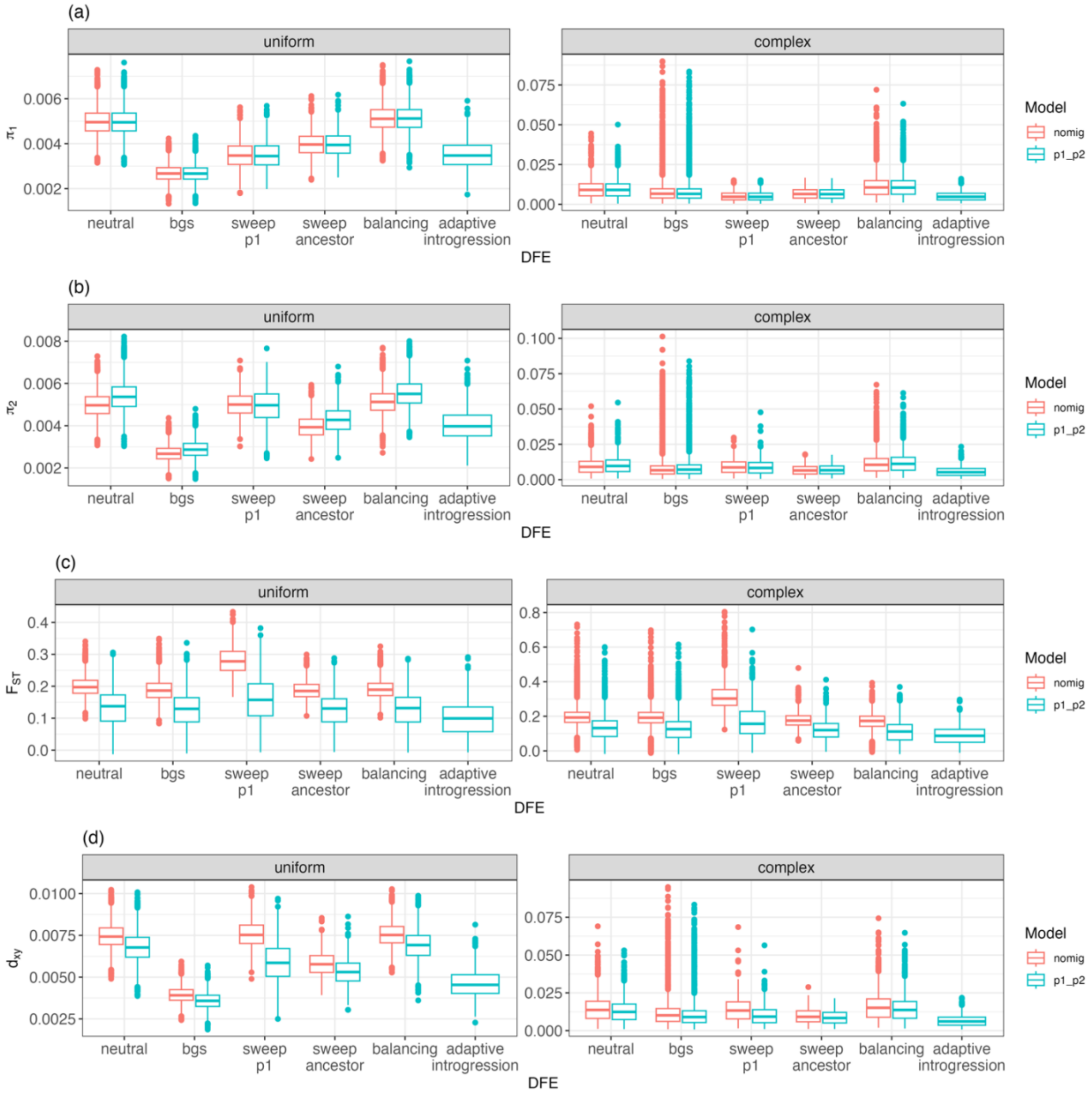


**Supporting Figure S1:** Summary statistics across models when T=1*N*. A) π within population 1; B) π within population 2; C) $F_{ST}$ between populations; D) $d_{xy}$ between populations.

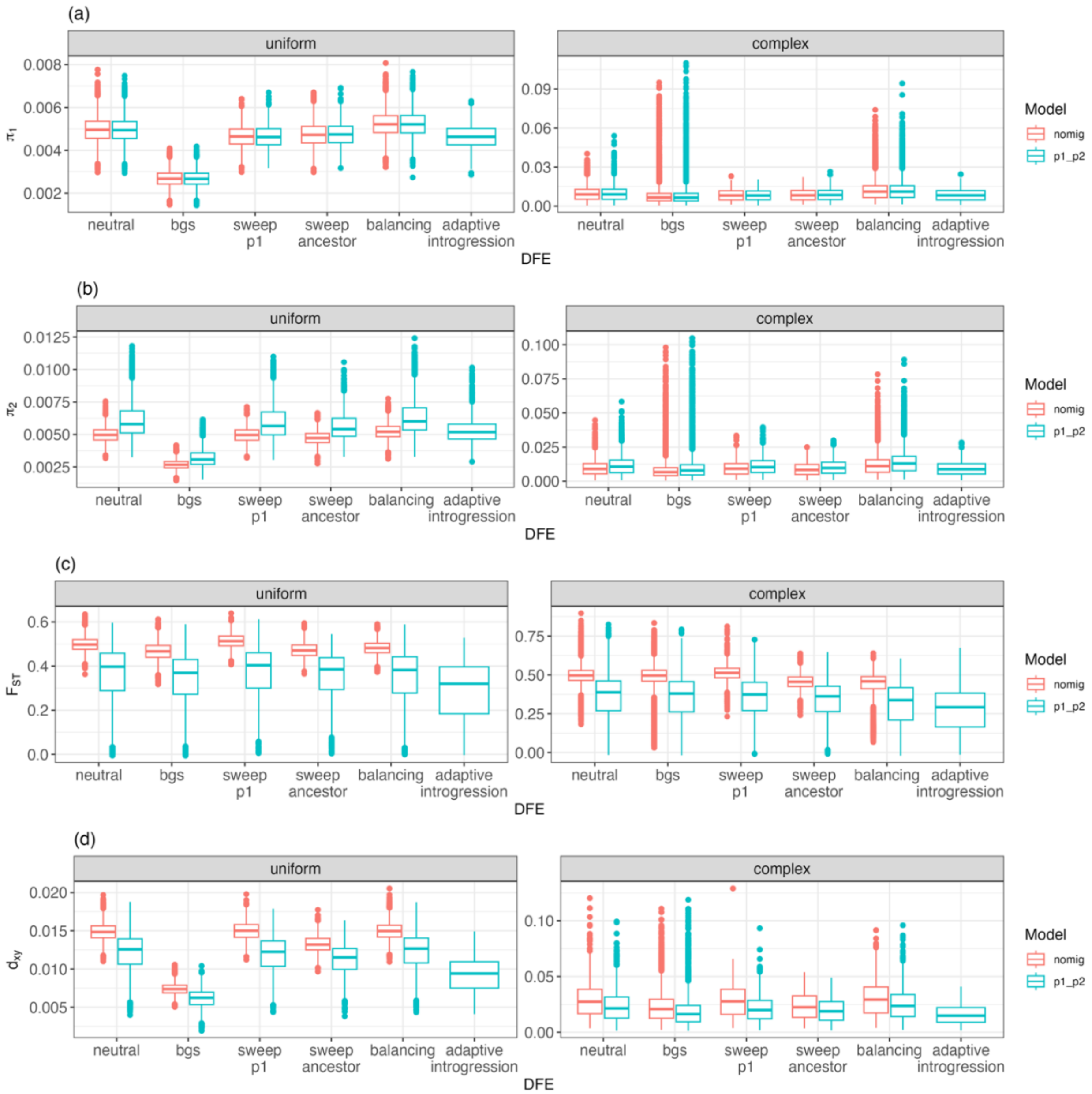


**Supporting Figure S2:** Summary statistics across models when T=4*N*. A) π within population 1; B) π within population 2; C) $F_{ST}$ between populations; D) $d_{xy}$ between populations.

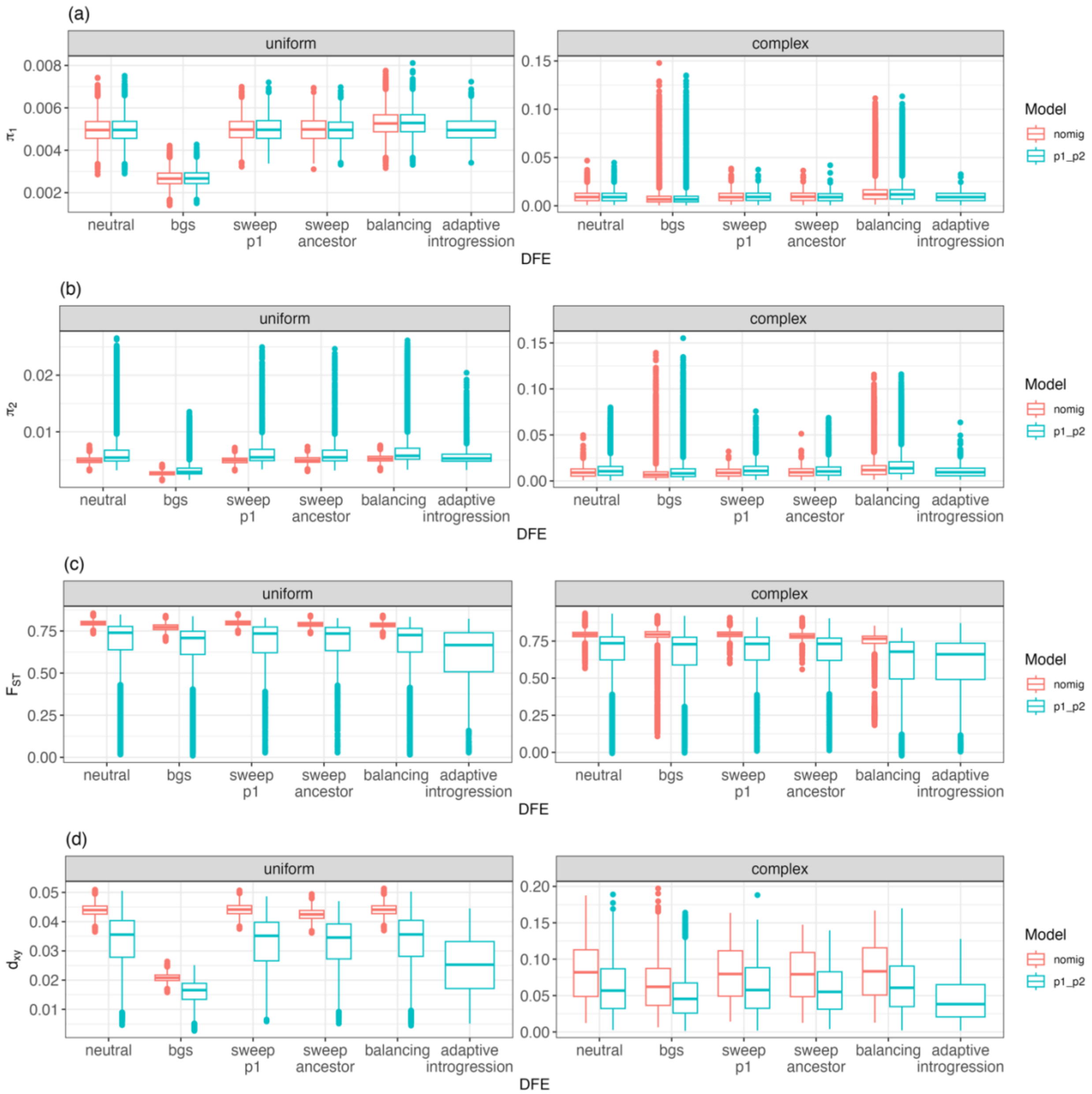


**Supporting Figure S3:** Summary statistics across models when T=16*N*. A) $\pi$ within population 1; B) $\pi$ within population 2; C) $F_{ST}$ between populations; D) $d_{xy}$ between populations.

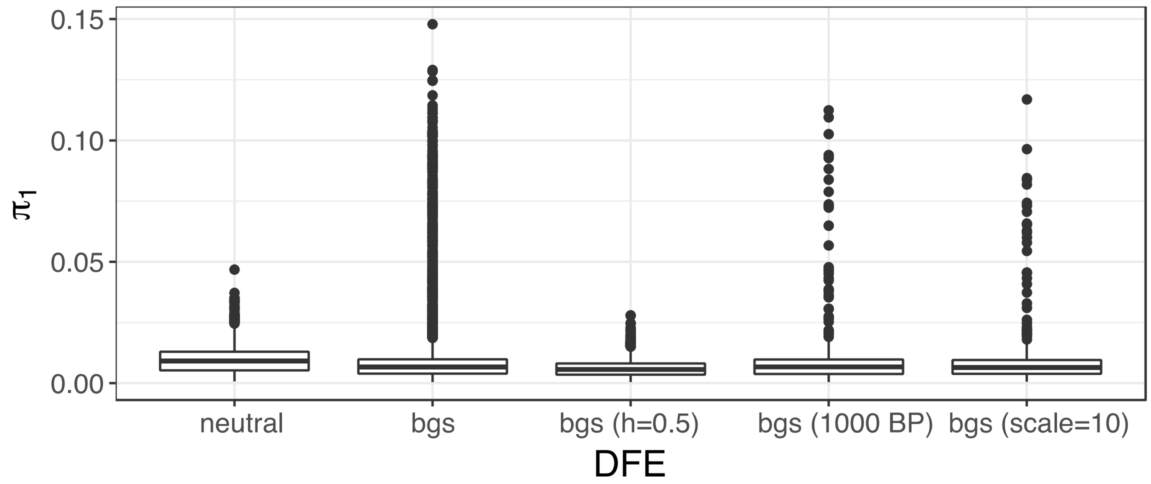


**Supporting Figure S4**. π within population 1 under the complex model when T=16*N* for the neutral case, the BGS case, a model with BGS and the dominance coefficient set to 0.5, the BGS case with 1000 randomly sampled replicates, and the BGS case with the scaling parameter set to 10 instead of 100.

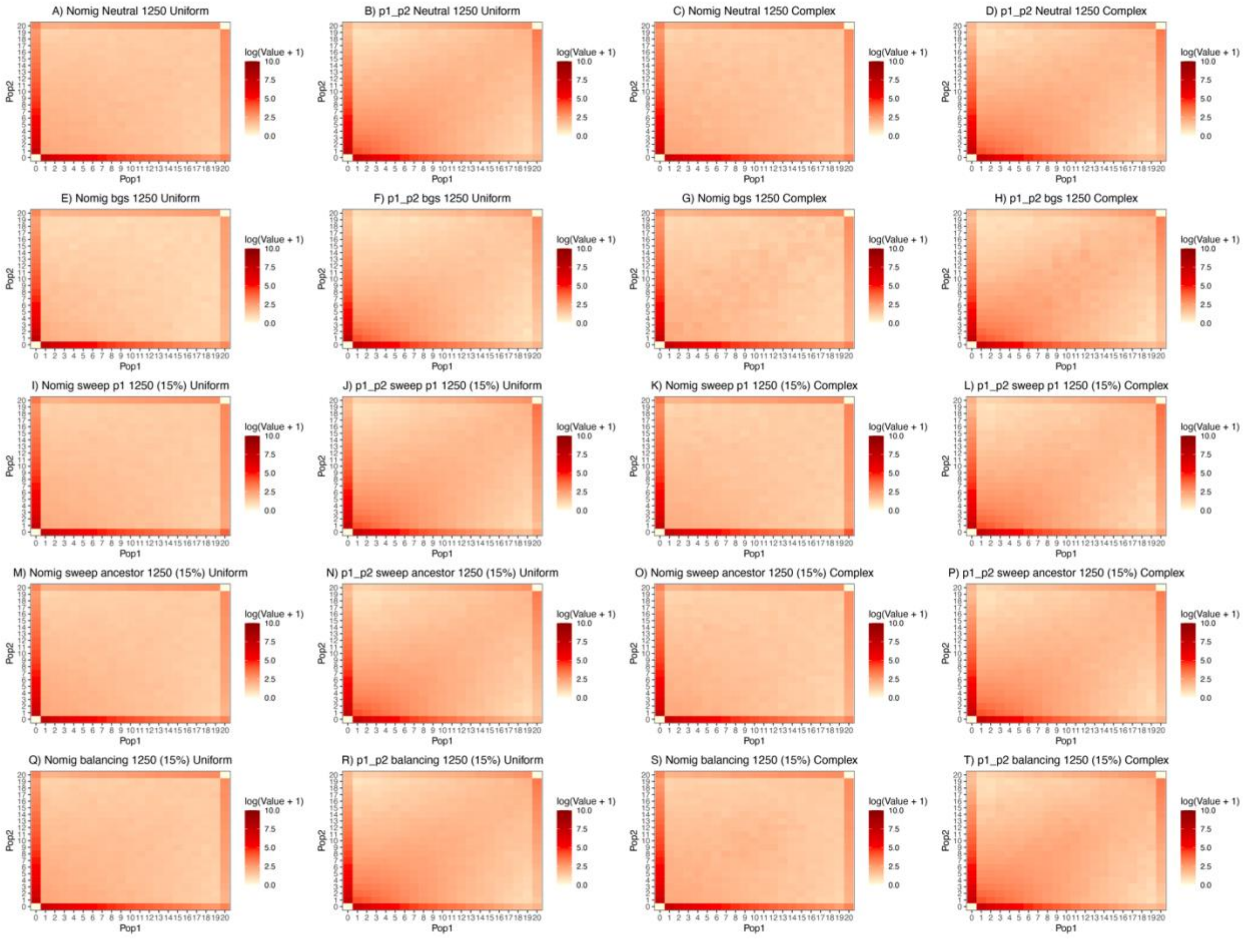


**Supporting Figure S5:** SFS under each model and condition in the when T=1*N*.

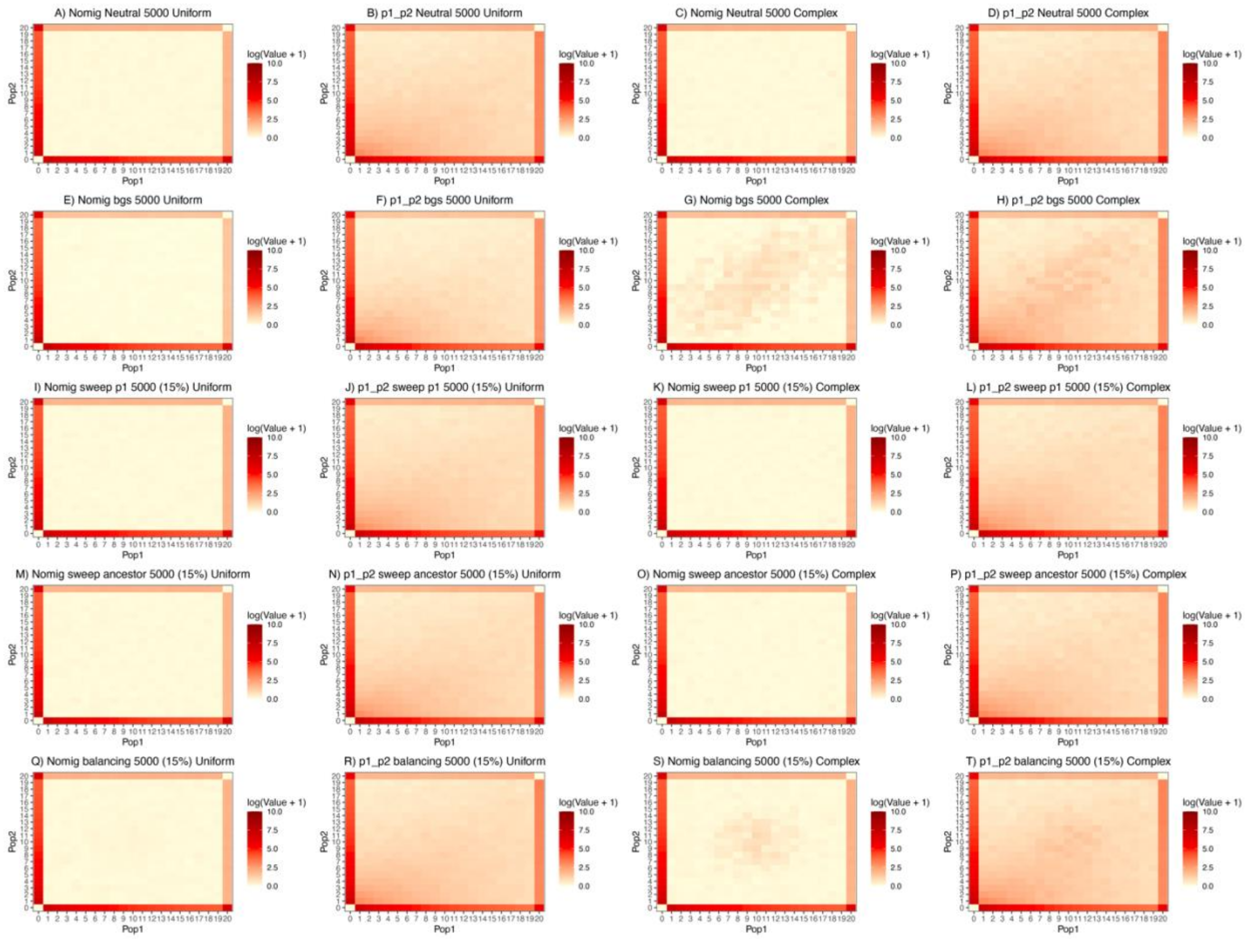

**Supporting Figure S6:** SFS under each model and condition when T=4*N*.

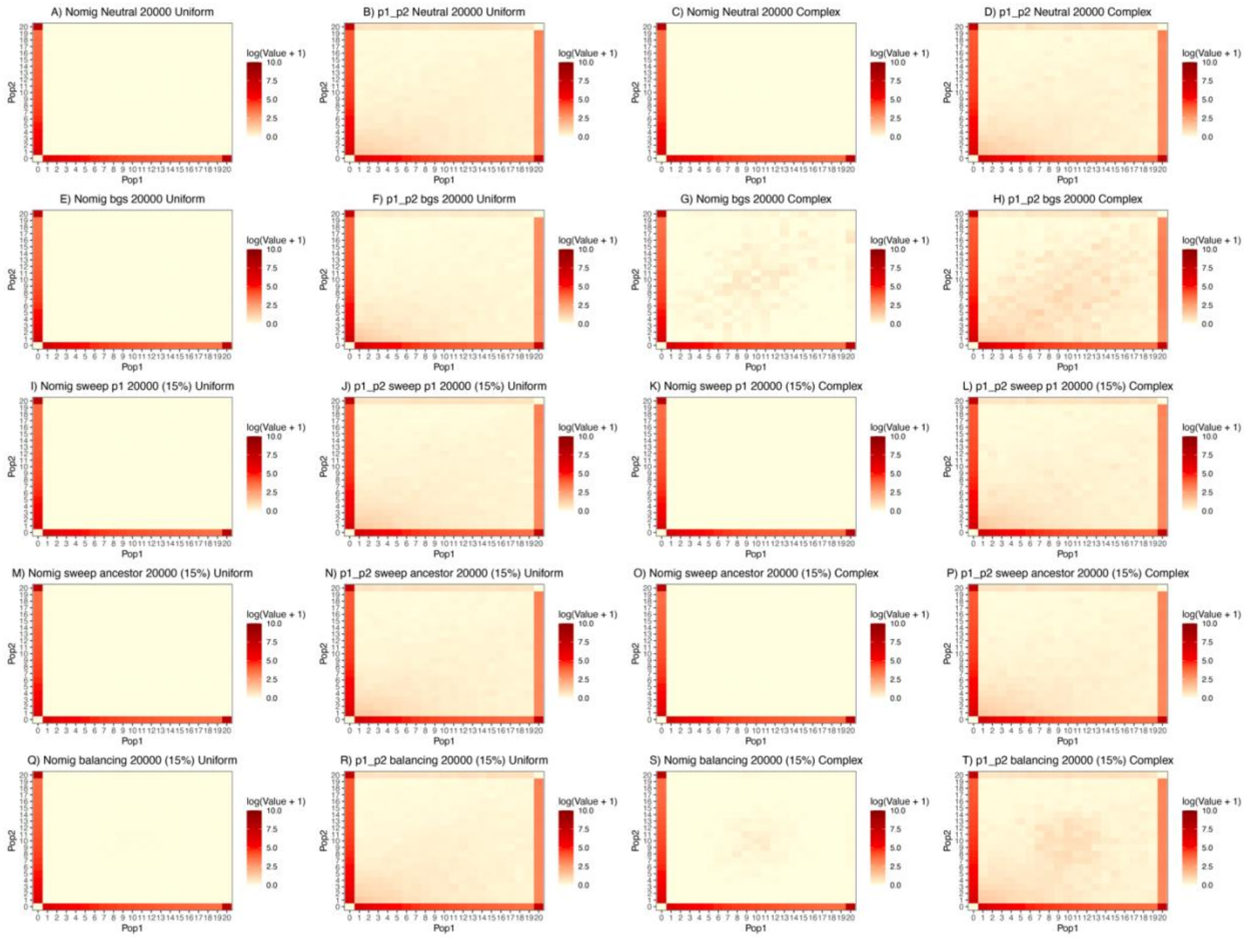


**Supporting Figure S7:** SFS under each model and condition when T=16*N*.

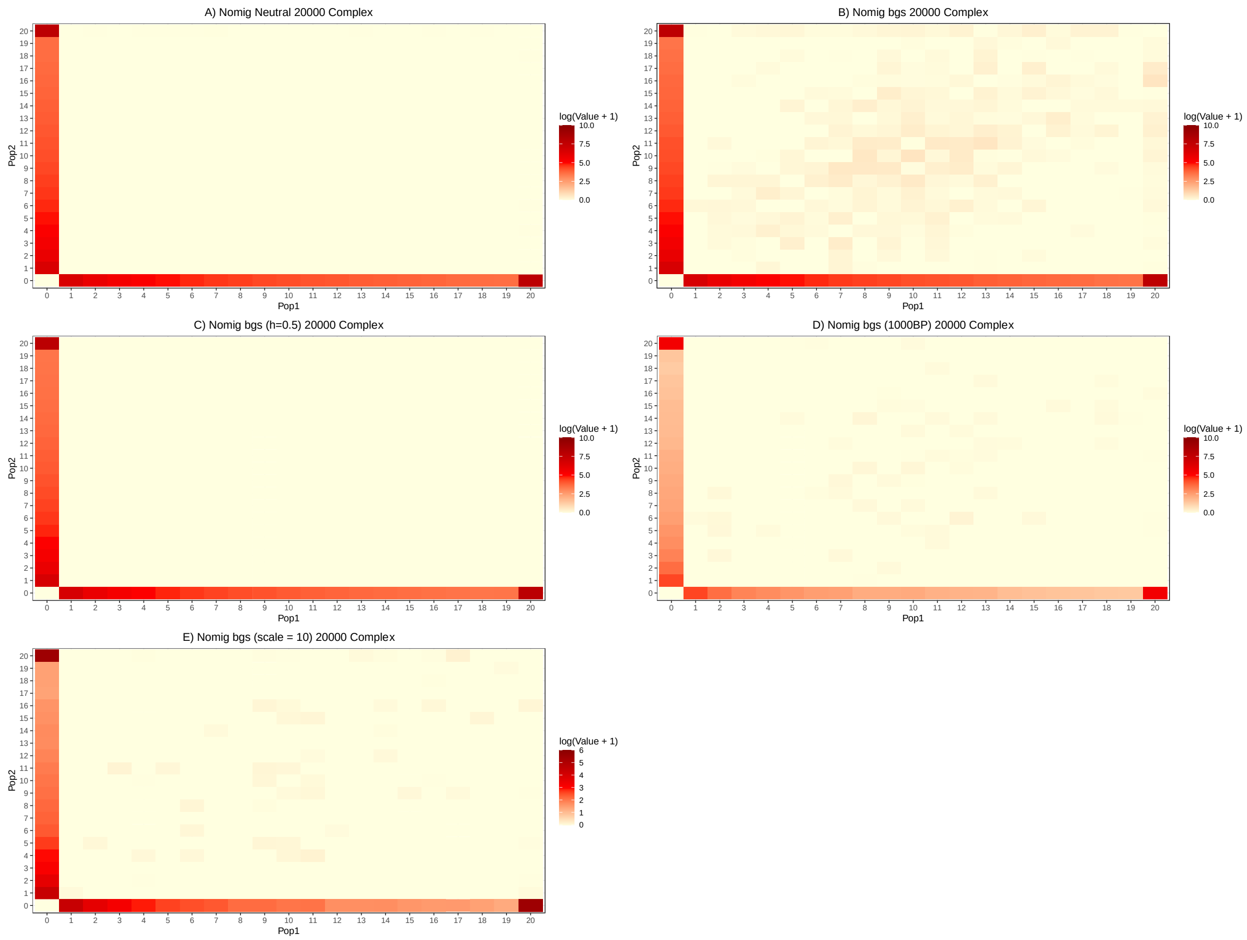


**Supporting Figure S8:** A comparison of SFS in the absence of migration with a complex genomic architecture when T=16*N* under neutral (A), BGS (B), BGS with the dominance coefficient set to 0.5 (C), BGS with 1000 bp (D), and BGS with 1000 bp scaled by 10 instead of 100 (E).

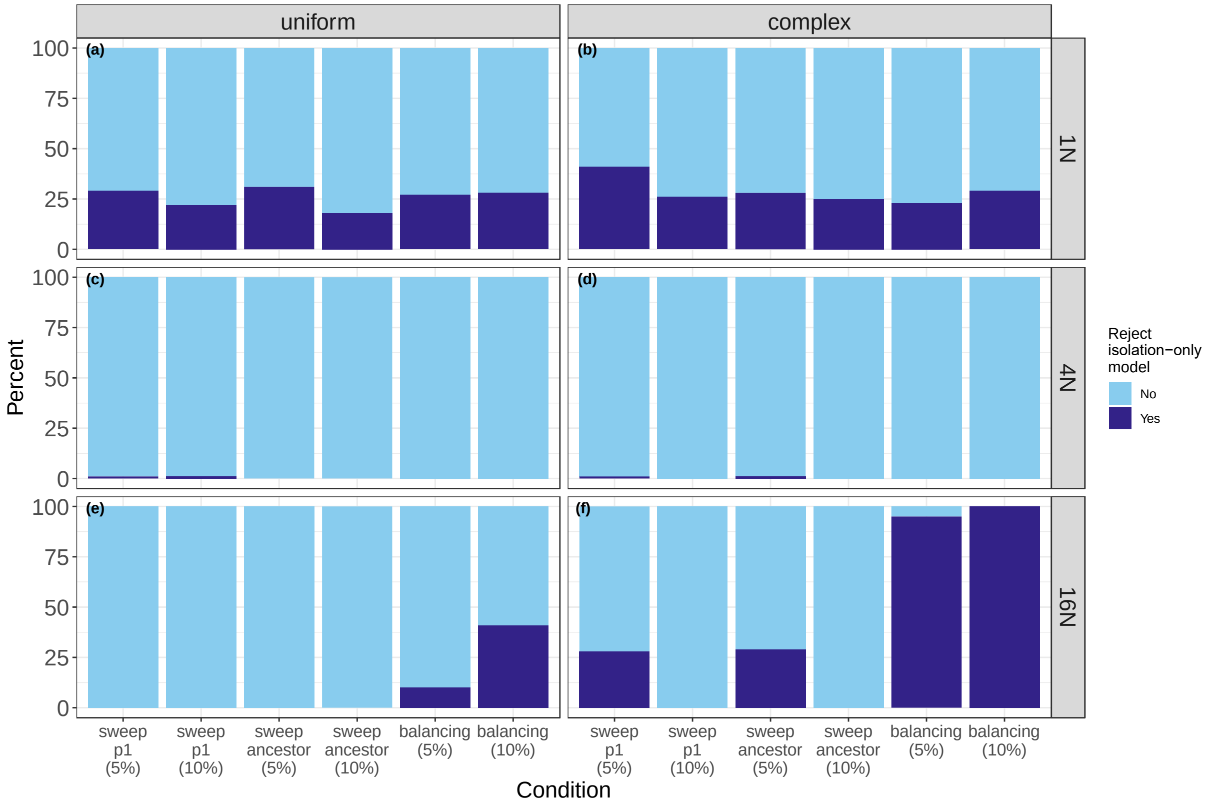


**Supporting Figure S9.** Results of the LRT in ∂a∂i when sweeps composed 5 or 10 percent of the data. Light blue indicates cases where we failed to reject the isolation-only model, and dark blue indicates cases where we rejected the isolation-only model in favor of the isolation-with-migration model. a) results for T=1*N* with a uniform genomic architecture; b) results for T=1*N* with a complex genomic architecture; c) results for T=4*N* with a uniform genomic architecture; d) results for T=4*N* with a complex genomic architecture; e) results for T=16*N* with a uniform genomic architecture; f) results for T=16*N* with a complex genomic architecture.

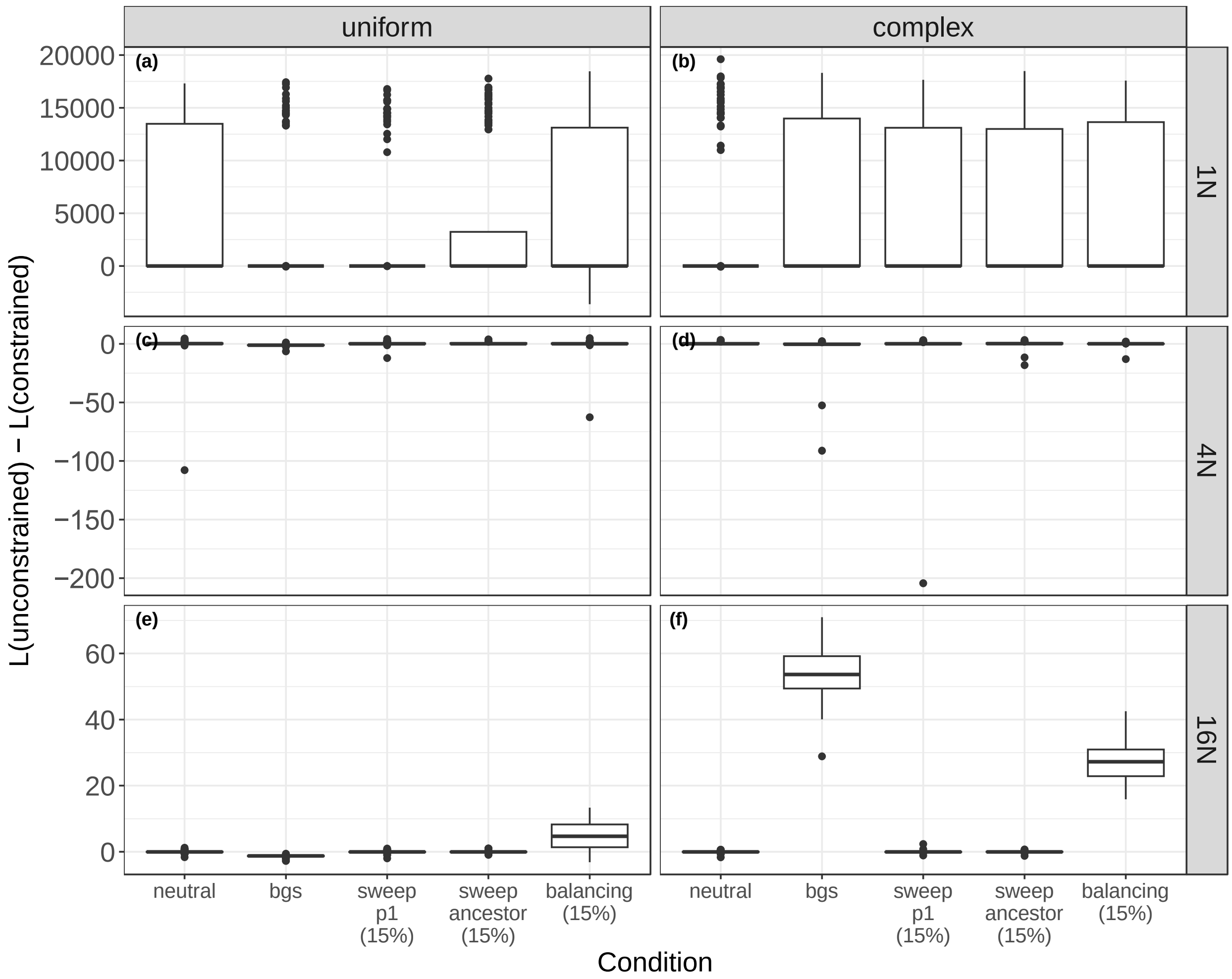


**Supporting Figure S10.** Difference in likelihoods between the unconstrained and constrained models on datasets without migration in ∂a∂i. a) results for T=1*N* with a uniform genomic architecture; b) results for T=1*N* with a complex genomic architecture; c) results for T=4*N* with a uniform genomic architecture; d) results for T=4*N* with a complex genomic architecture; e) results for T=16*N* with a uniform genomic architecture; f) results for T=16*N* with a complex genomic architecture.

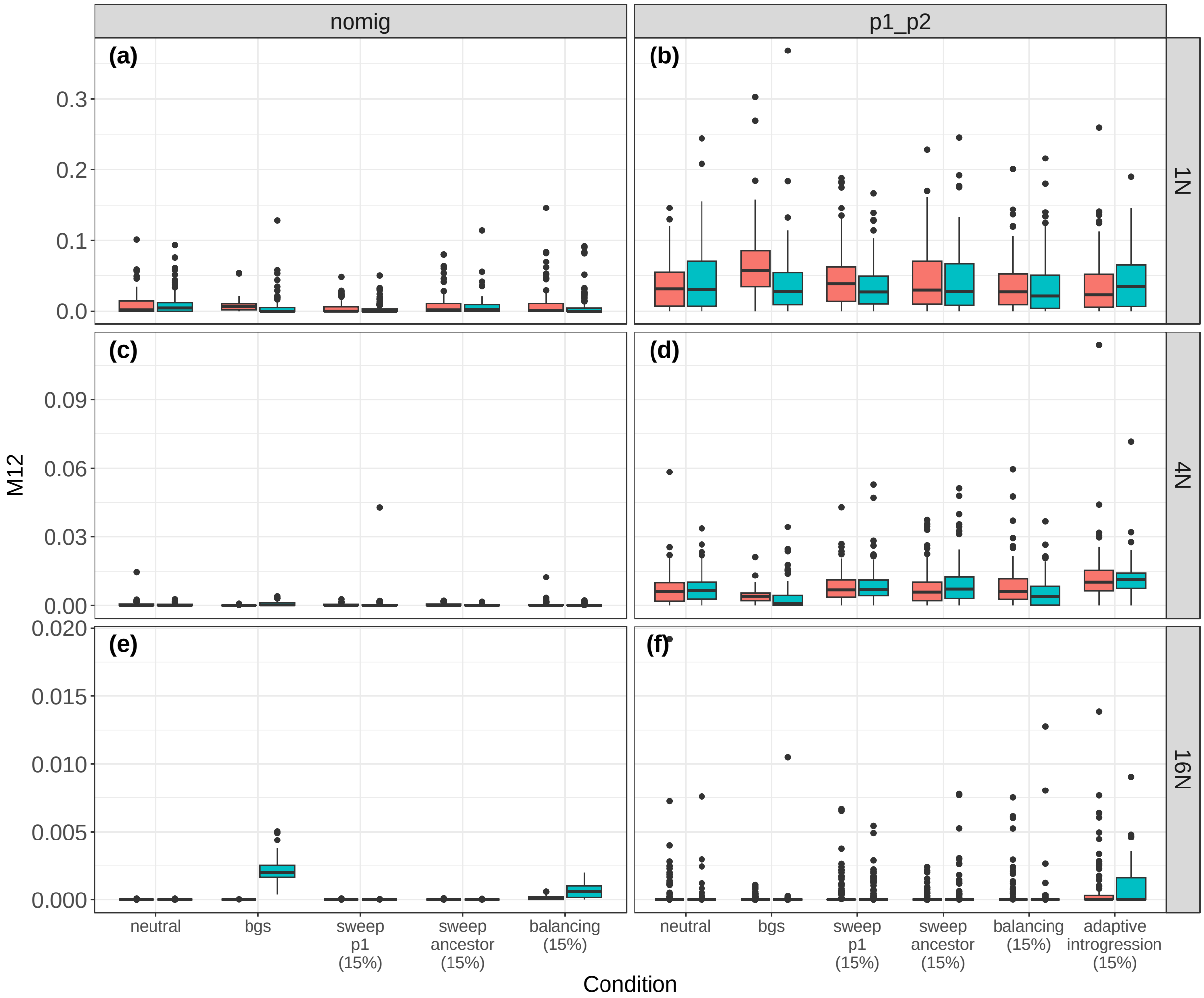


**Supporting Figure S11.** Estimates of $M_{12}$ in ∂a∂i. Estimates are in units of $2 \times N_{ref} \times m_{ij}$.The colors indicate the uniform (pink) and complex (blue) genomic architectures. a) results for T=1*N* and the nomig model; b) results for T=1*N* and the p1_p2 model; c) results for T=4*N* and the nomig model; d) results for T=4*N* and the p1_p2 model; e) results for T=16*N* and the nomig model; f) results for T=16*N* and the p1_p2 model.

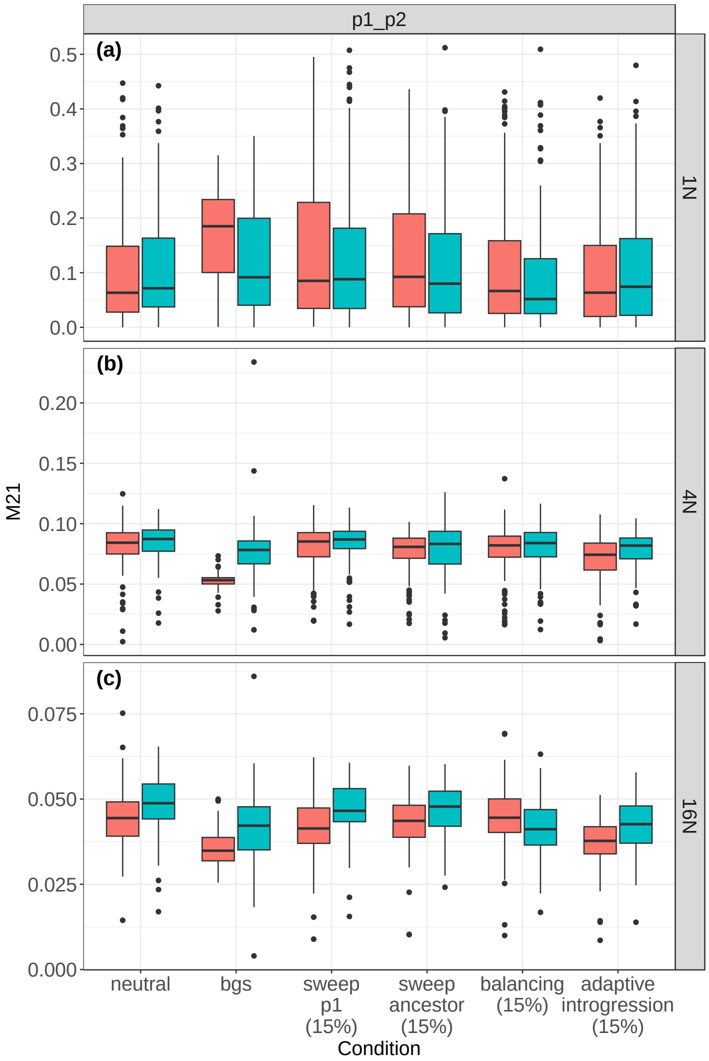


**Supporting Figure S12.** Estimates of $M_{21}$ in ∂a∂i for models including migration. Estimates are in units of $2 \times N_{ref} \times m_{ij}$. The colors indicate the uniform (pink) and complex (blue) genomic architectures. a) results for T=1*N* and the p1_p2 model; b) results for T=4*N* and the p1_p2 model; c) results for T=16*N* and the p1_p2 model.

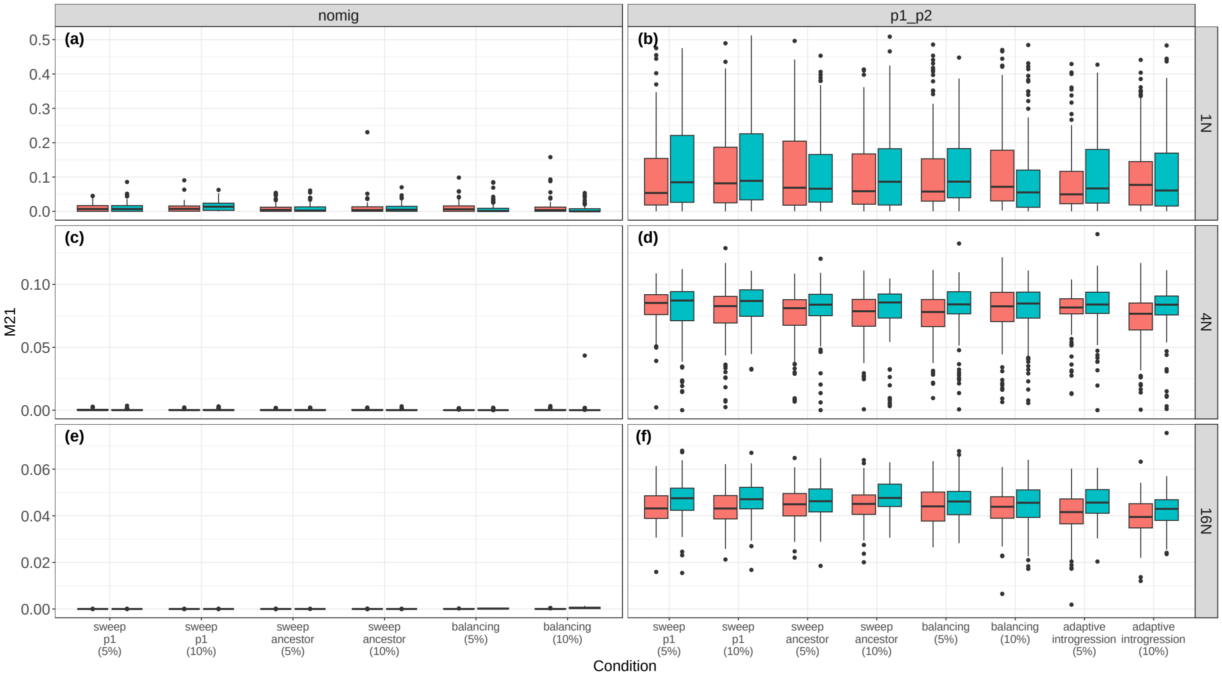


**Supporting Figure S13.** Estimates of $M_{21}$ in ∂a∂i with different percentages of loci experiencing selective sweeps. Estimates are in units of $2 \times N_{ref} \times m_{ij}$. The colors indicate the uniform (pink) and complex (blue) genomic architectures. a) results for T=1*N* and the nomig model; b) results for T=1*N* and the p1_p2 model; c) results for T=4*N* and the nomig model; d) results for T=4*N* and the p1_p2 model; e) results for T=16*N* and the nomig model; f) results for T=16*N* and the p1_p2 model.

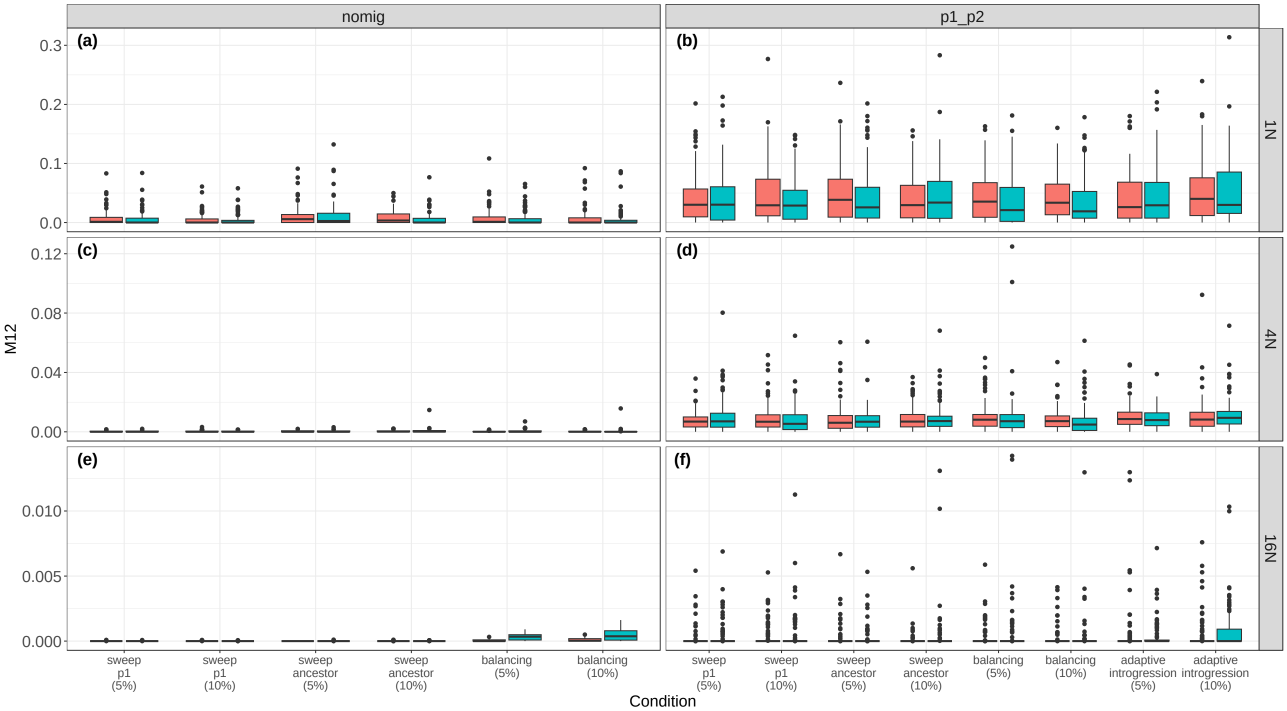


**Supporting Figure S14.** Estimates of $M_{12}$ in ∂a∂i with different percentages of loci experiencing selective sweeps. Estimates are in units of $2 \times N_{ref} \times m_{ij}$. The colors indicate the uniform (pink) and complex (blue) genomic architectures. a) results for T=1*N* and the nomig model; b) results for T=1*N* and the p1_p2 model; c) results for T=4*N* and the nomig model; d) results for T=4*N* and the p1_p2 model; e) results for T=16*N* and the nomig model; f) results for T=16*N* and the p1_p2 model.

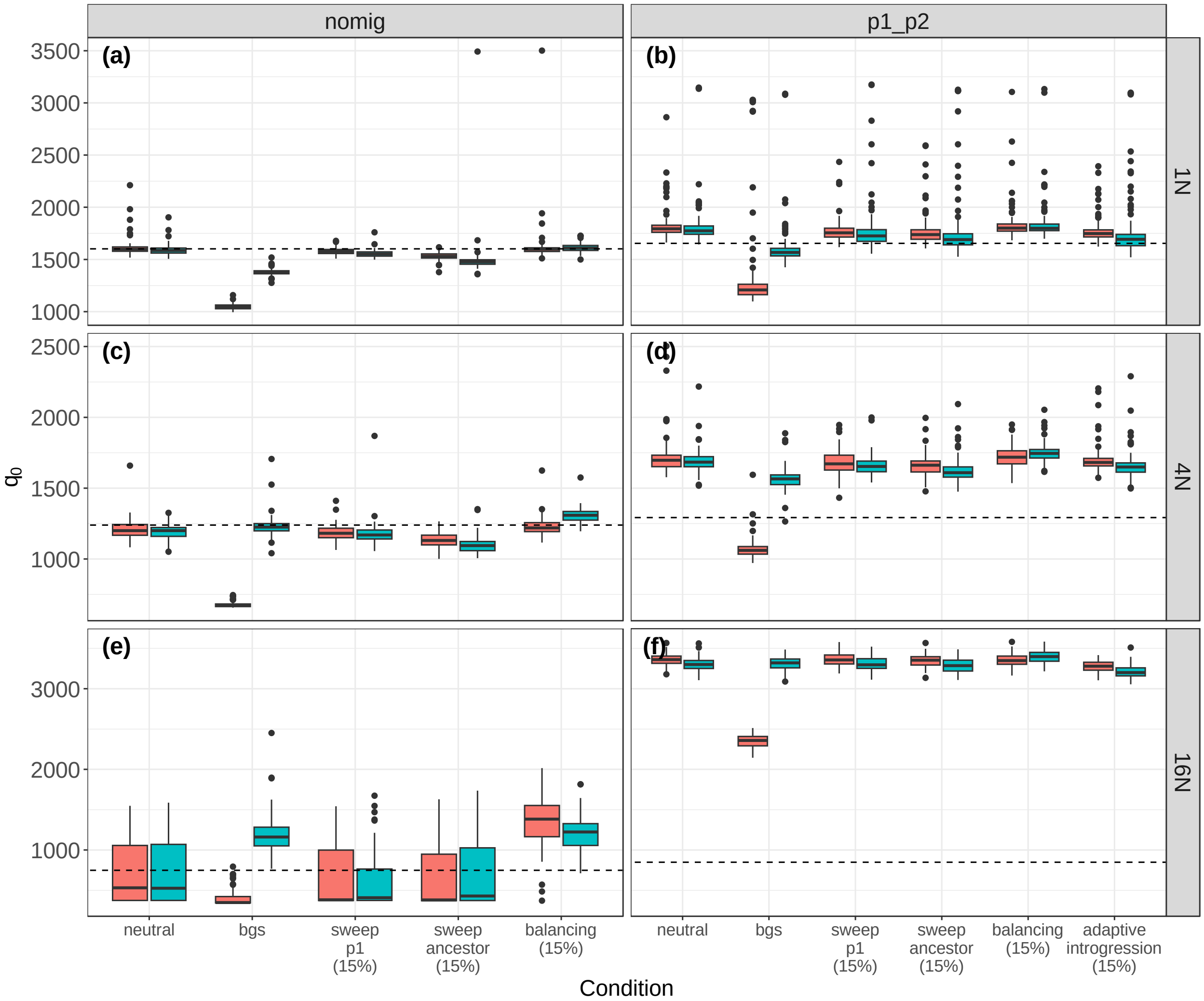


**Supporting Figure S15.** Estimates of $\theta_0$ in ∂a∂i. Estimates are in units of $\theta = 4 \times N_{ref} \times \mu \times L$, where L is the number of sites used to construct the SFS. The colors indicate the uniform (pink) and complex (blue) genomic architectures. a) results for T=1*N* and the nomig model; b) results for T=1*N* and the p1_p2 model; c) results for T=4*N* and the nomig model; d) results for T=4*N* and the p1_p2 model; e) results for T=16*N* and the nomig model; f) results for T=16*N* and the p1_p2 model.

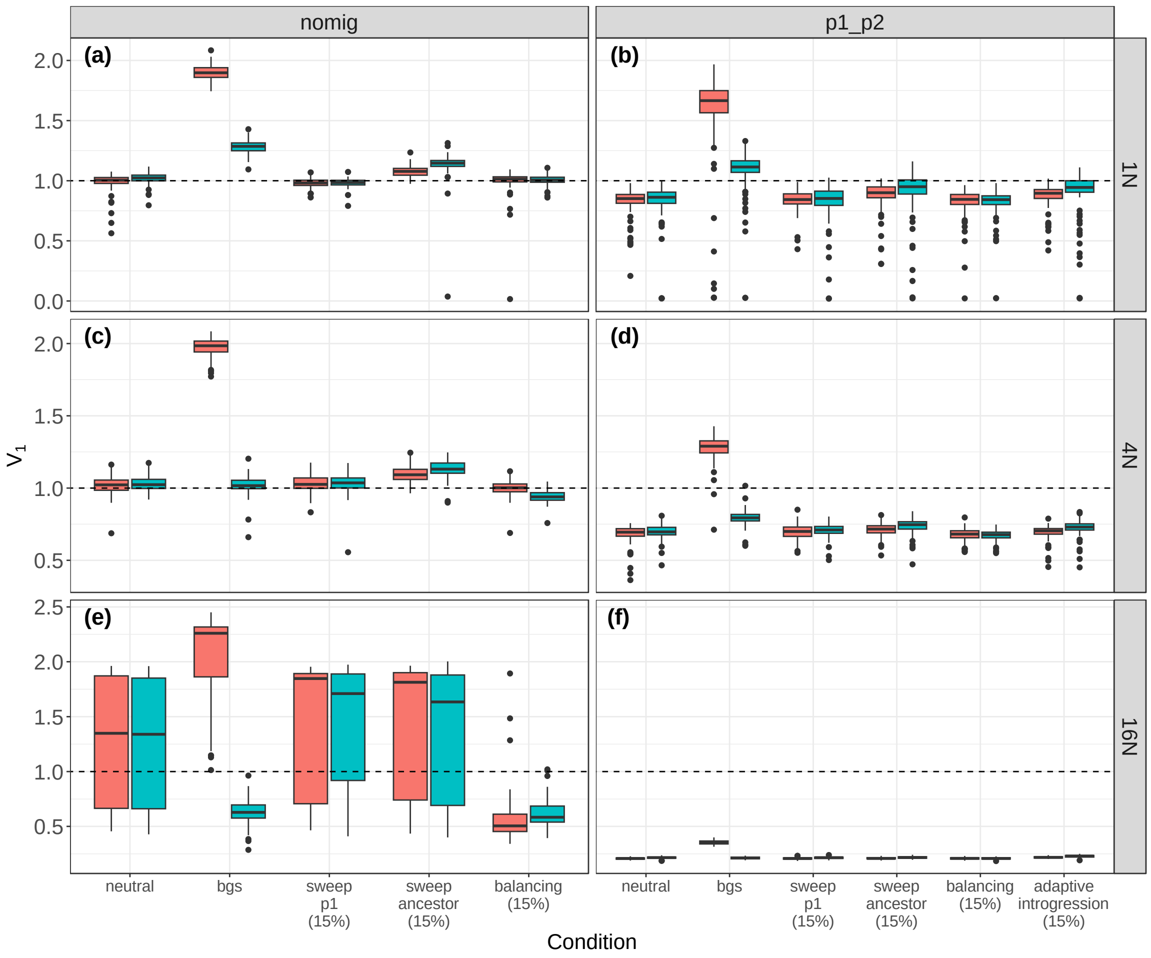


**Supporting Figure S16.** Estimates of $V_1$ in ∂a∂i. $V_1$ is the size of population 1 relative to the ancestral population. The colors indicate the uniform (pink) and complex (blue) genomic architectures. a) results for T=1*N* and the nomig model; b) results for T=1*N* and the p1_p2 model; c) results for T=4*N* and the nomig model; d) results for T=4*N* and the p1_p2 model; e) results for T=16*N* and the nomig model; f) results for T=16*N* and the p1_p2 model.

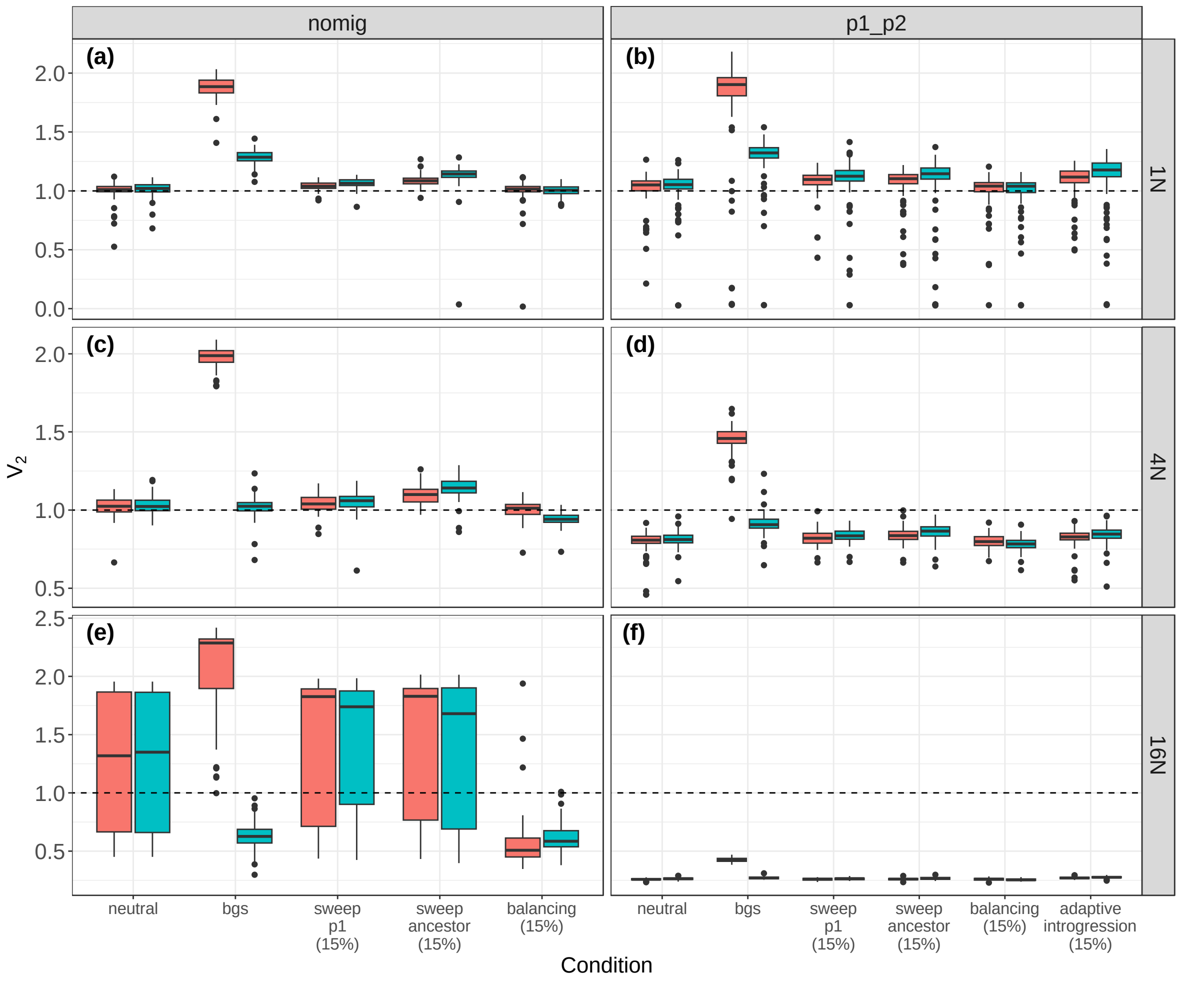


**Supporting Figure S17.** Estimates of $V_2$ in ∂a∂i. $V_2$ is the size of population 2 relative to the ancestral population. The colors indicate the uniform (pink) and complex (blue) genomic architectures. a) results for T=1*N* and the nomig model; b) results for T=1*N* and the p1_p2 model; c) results for T=4*N* and the nomig model; d) results for T=4*N* and the p1_p2 model; e) results for T=16*N* and the nomig model; f) results for T=16*N* and the p1_p2 model.

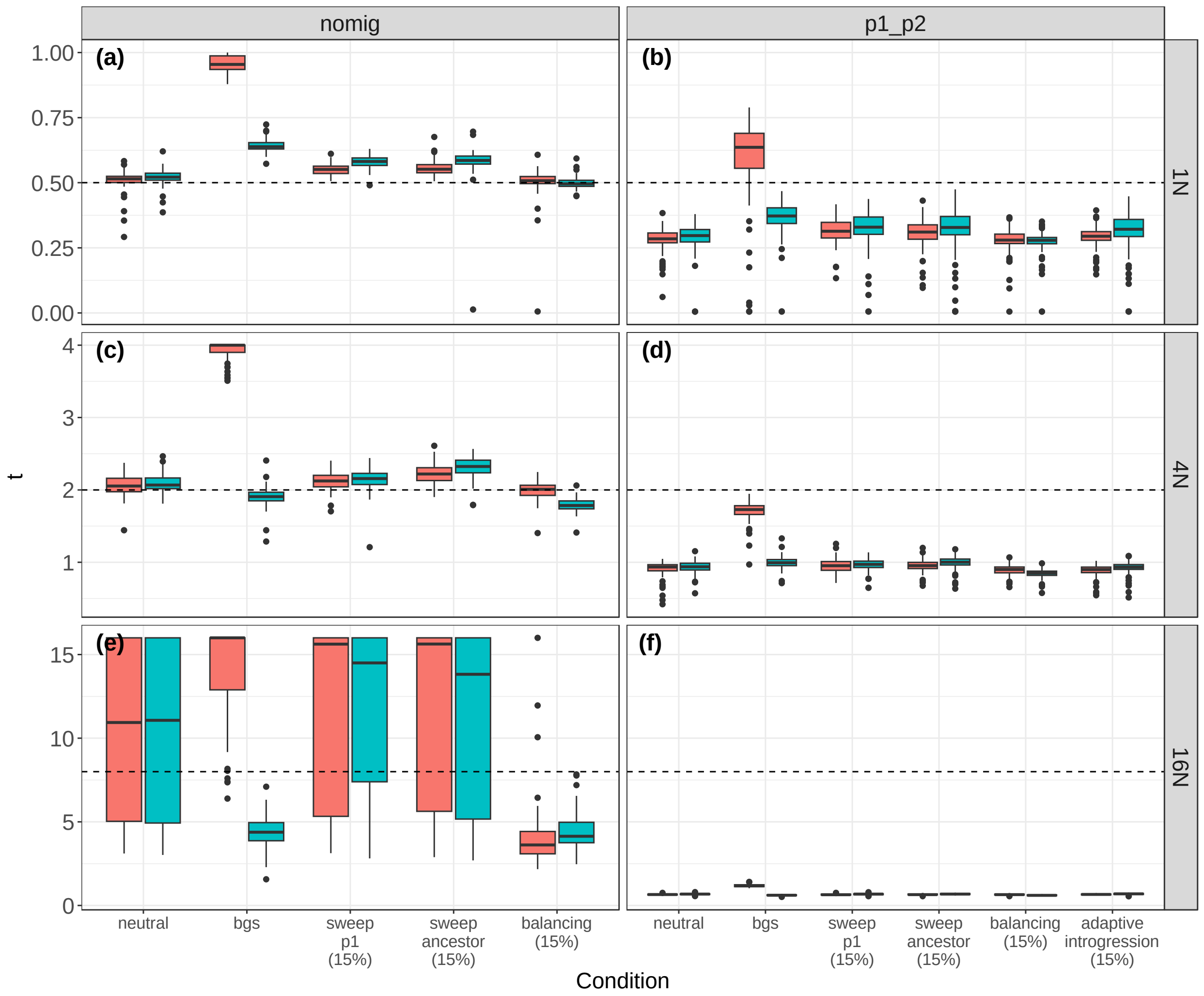


**Supporting Figure S18.** Estimates of τ in ∂a∂i. Estimates are in units of $2N_{ref}$ generations. The colors indicate the uniform (pink) and complex (blue) genomic architectures. a) results for T=1*N* and the nomig model; b) results for T=1*N* and the p1_p2 model; c) results for T=4*N* and the nomig model; d) results for T=4*N* and the p1_p2 model; e) results for T=16*N* and the nomig model; f) results for T=16*N* and the p1_p2 model.

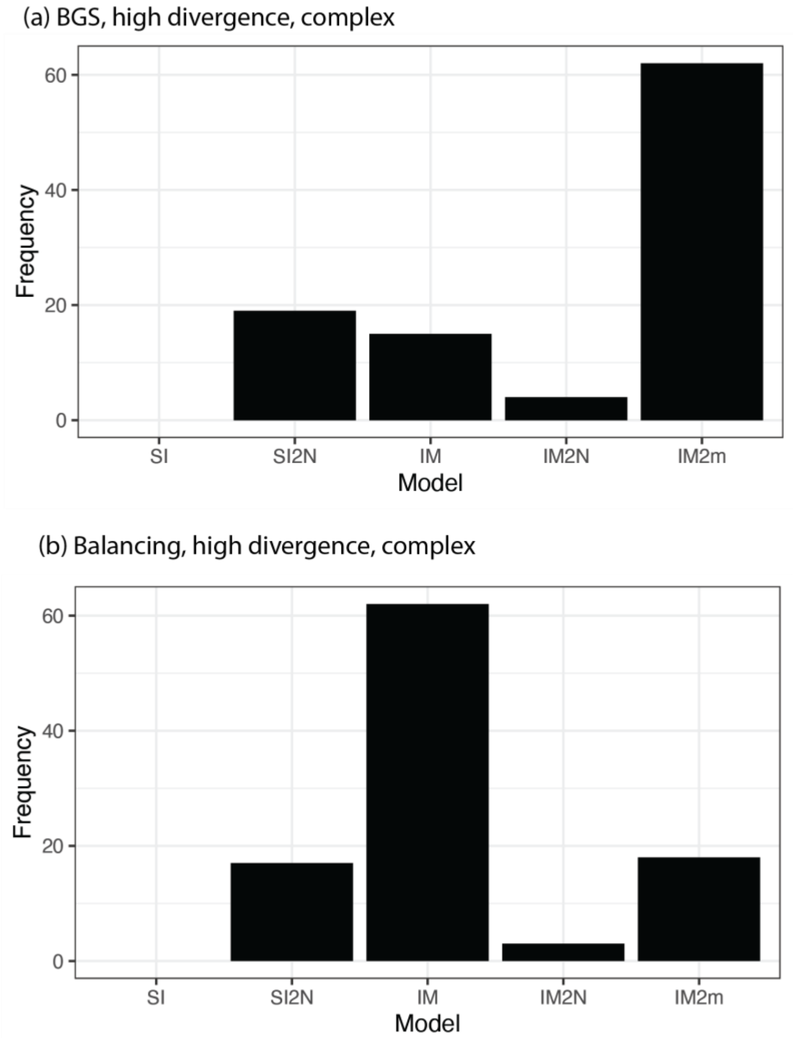


**Supporting Figure S19.** Results from the approach used by Rougeux *et al.* (2017) to accommodate selection in ∂a∂i. (a) Results with BGS, T=16*N*, and a complex genomic architecture. (b) Results with balancing selection, T=16*N*, and a complex genomic architecture.

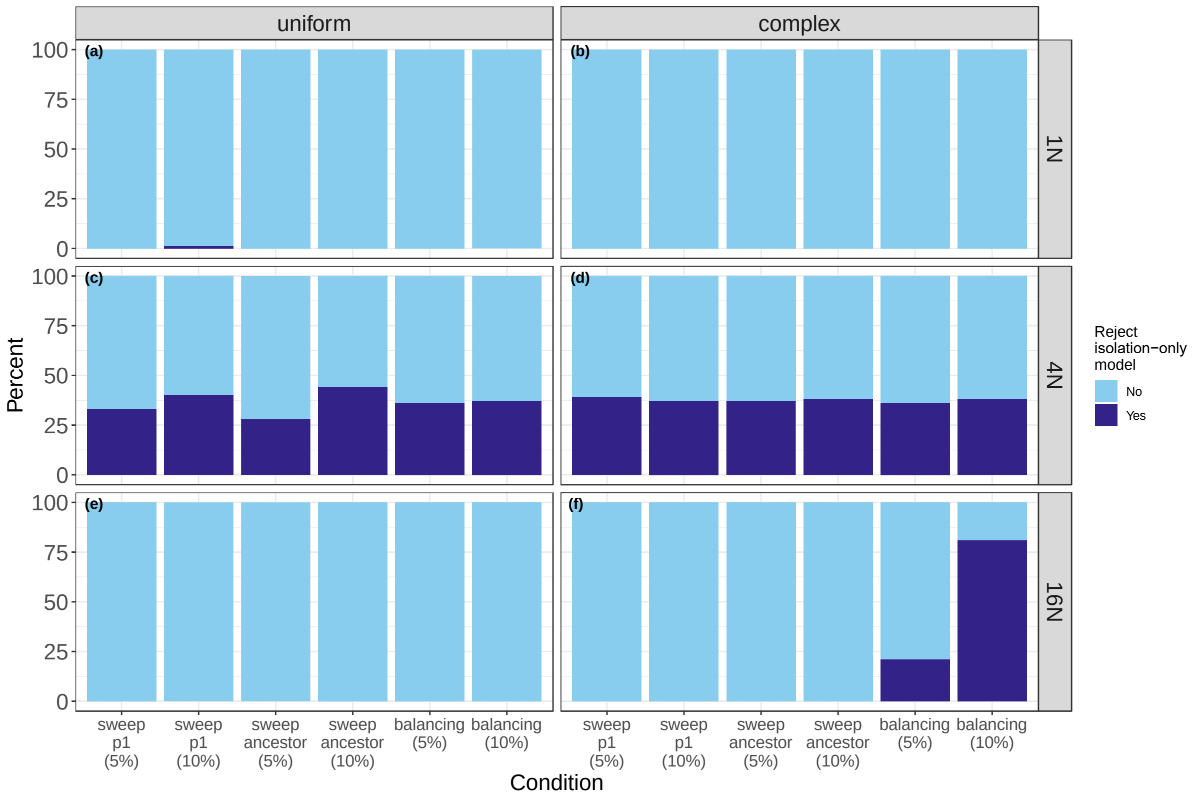


**Supporting Figure S20.** Results of the LRT in fastsimcoal2 when sweeps composed 5 or 10 percent of the data. Light blue indicates cases where we failed to reject the isolation-only model, and dark blue indicates cases where we rejected the isolation-only model in favor of the isolation-with-migration model. a) results for T=1*N* with a uniform genomic architecture; b) results for T=1*N* with a complex genomic architecture; c) results for T=4*N* with a uniform genomic architecture; d) results for T=4*N* with a complex genomic architecture; e) results for T=16*N* with a uniform genomic architecture; f) results for T=16*N* with a complex genomic architecture.

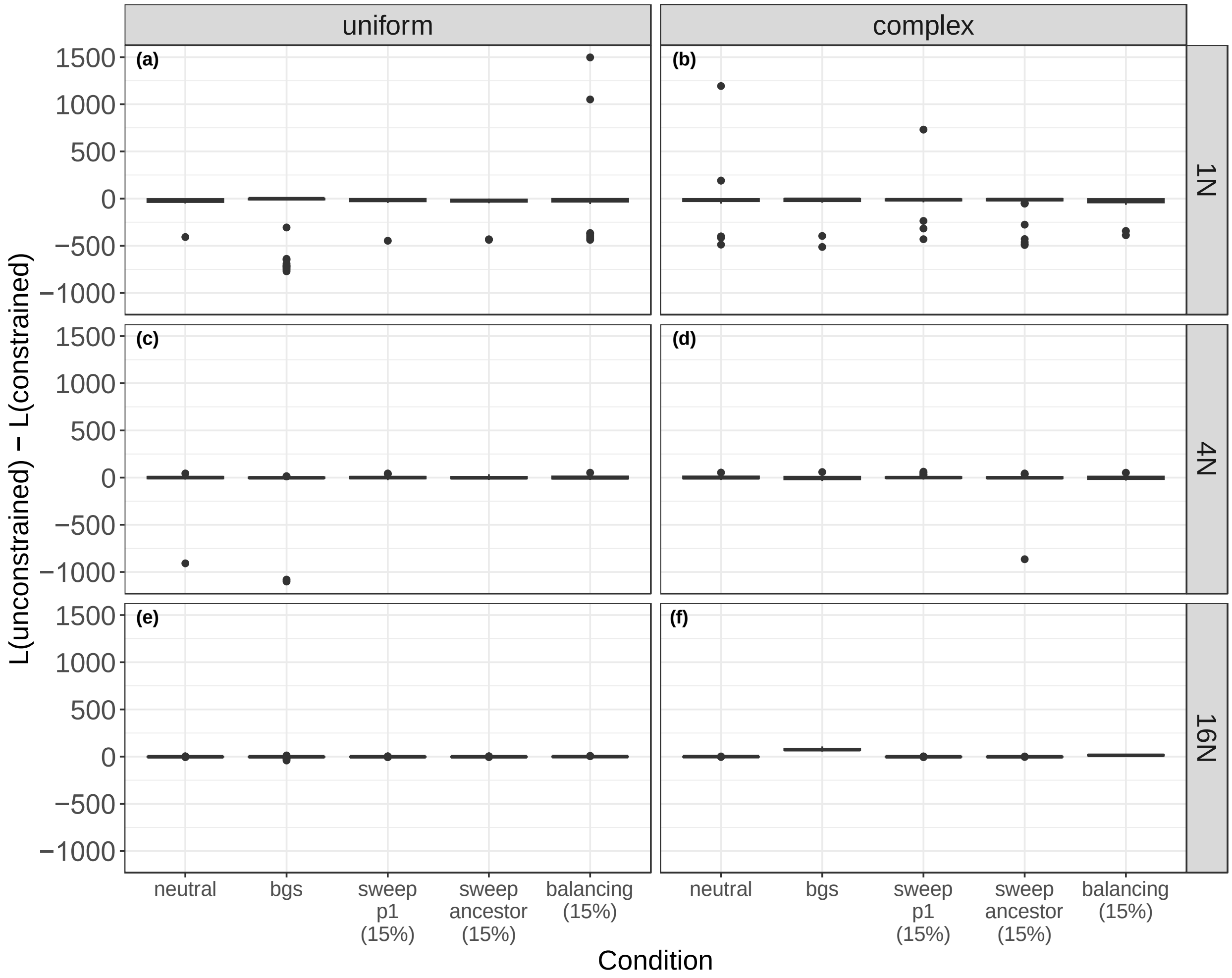


**Supporting Figure S21.** Difference in likelihoods between the unconstrained and constrained models on datasets without migration in fastsimcoal2. a) results for T=1*N* with a uniform genomic architecture; b) results for T=1*N* with a complex genomic architecture; c) results for T=4*N* with a uniform genomic architecture; d) results for T=4*N* with a complex genomic architecture; e) results for T=16*N* with a uniform genomic architecture; f) results for T=16*N* with a complex genomic architecture.

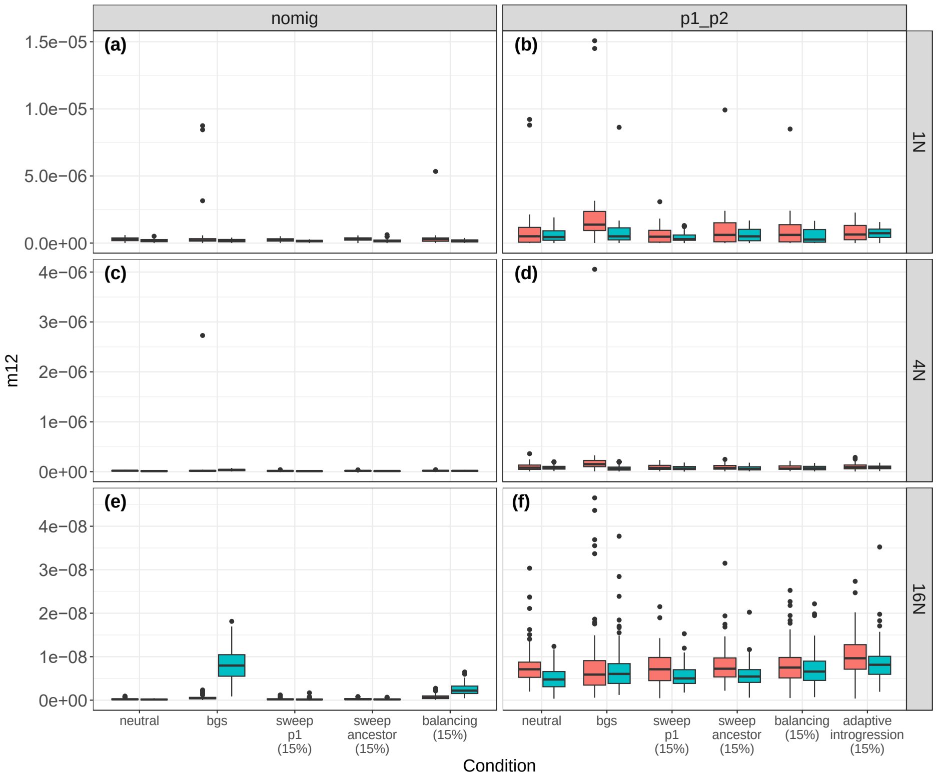


**Supporting Figure S22.** Estimates of $m_{12}$ in fastsimcoal2. The rate $m_{ij}$ is the probability of any gene moving from population *i* to population *j* backwards in time each generation. The colors indicate the uniform (pink) and complex (blue) genomic architectures. a) results for T=1*N* and the nomig model; b) results for T=1*N* and the p1_p2 model; c) results for T=4*N* and the nomig model; d) results for T=4*N* and the p1_p2 model; e) results for T=16*N* and the nomig model; f) results for T=16*N* and the p1_p2 model.

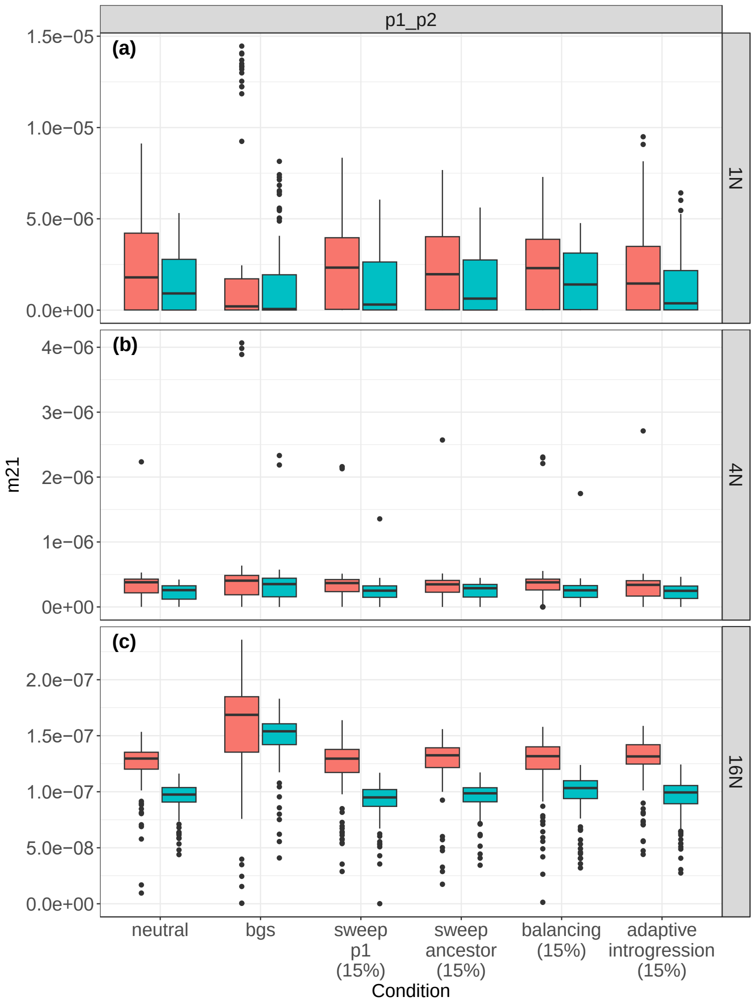


**Supporting Figure S23.** Estimates of $m_{21}$ in fastsimcoal2 for models including migration. The rate $m_{ij}$ is the probability of any gene moving from population $i$ to population $j$ backwards in time each generation. The colors indicate the uniform (pink) and complex (blue) genomic architectures. a) results for T=1*N* and the p1_p2 model; b) results for T=4*N* and the p1_p2 model; c) results for T=16*N* and the p1_p2 model.

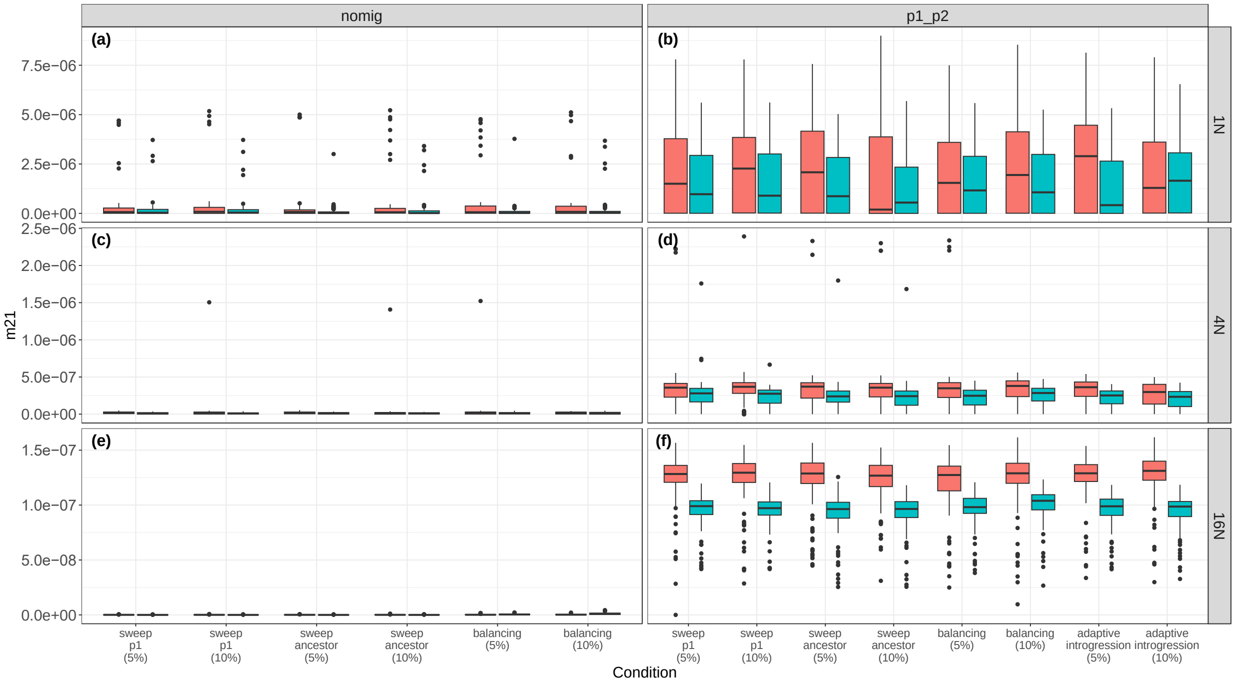


**Supporting Figure S24.** Estimates of $m_{21}$ in fastsimcoal2 with different percentages of loci experiencing selective sweeps. The rate $m_{ij}$ is the probability of any gene moving from population $i$ to population $j$ backwards in time each generation. The colors indicate the uniform (pink) and complex (blue) genomic architectures. a) results for T=1*N* and the nomig model; b) results for T=1*N* and the p1_p2 model; c) results for T=4*N* and the nomig model; d) results for T=4*N* and the p1_p2 model; e) results for T=16*N* and the nomig model; f) results for T=16*N* and the p1_p2 model.

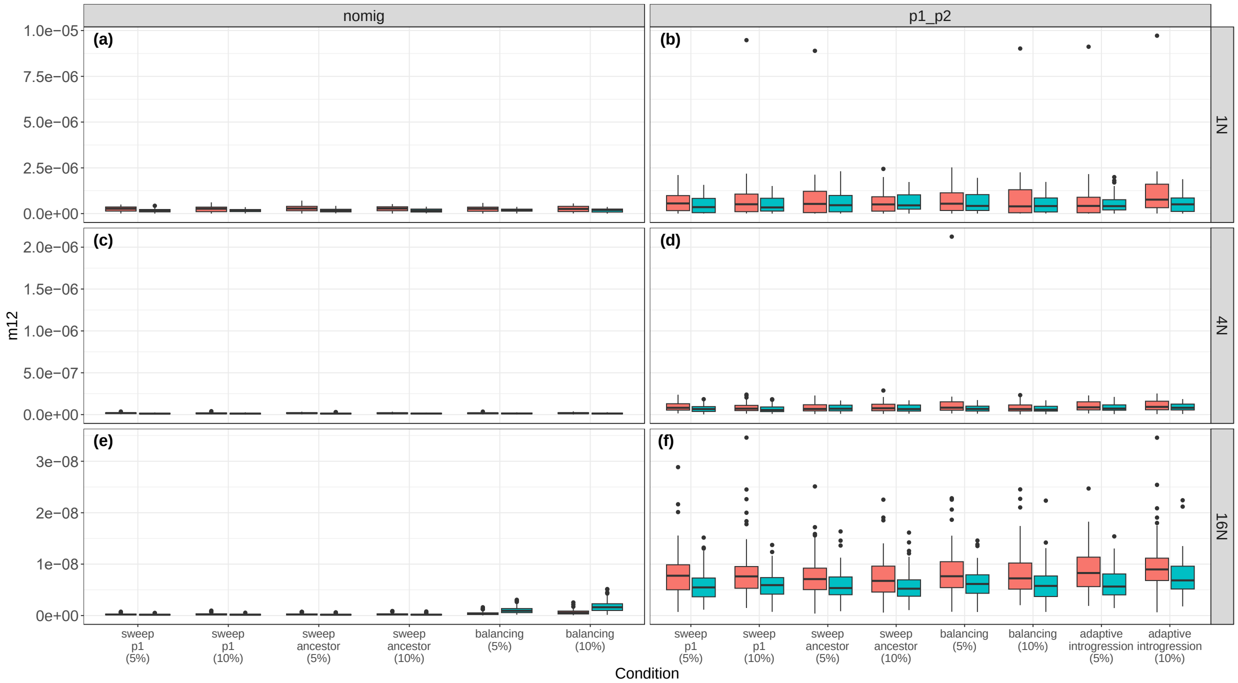


**Supporting Figure S25.** Estimates of $m_{12}$ in fastsimcoal2 with different percentages of loci experiencing selective sweeps. The rate $m_{ij}$ is the probability of any gene moving from population *i* to population *j* backwards in time each generation. The colors indicate the uniform (pink) and complex (blue) genomic architectures. a) results for T=1*N* and the nomig model; b) results for T=1*N* and the p1_p2 model; c) results for T=4*N* and the nomig model; d) results for T=4*N* and the p1_p2 model; e) results for T=16*N* and the nomig model; f) results for T=16*N* and the p1_p2 model.

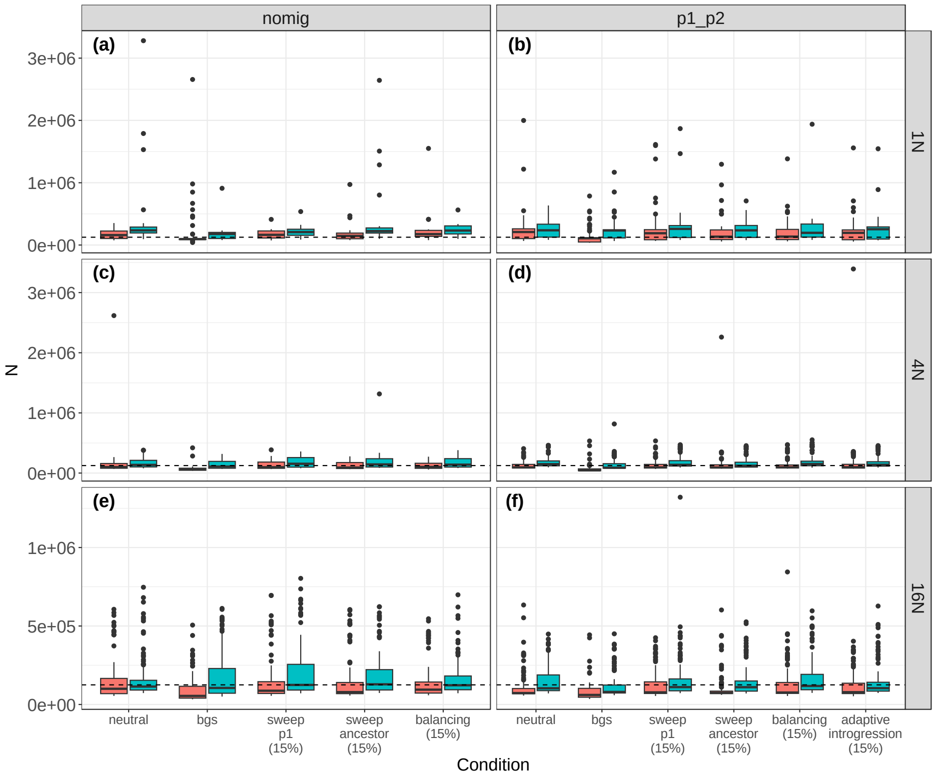


**Supporting Figure S26.** Estimates of $N_{ANC}$ in fastsimcoal2. The colors indicate the uniform (pink) and complex (blue) genomic architectures. a) results for T=1*N* and the nomig model; b) results for T=1*N* and the p1_p2 model; c) results for T=4*N* and the nomig model; d) results for T=4*N* and the p1_p2 model; e) results for T=16*N* and the nomig model; f) results for T=16*N* and the p1_p2 model.

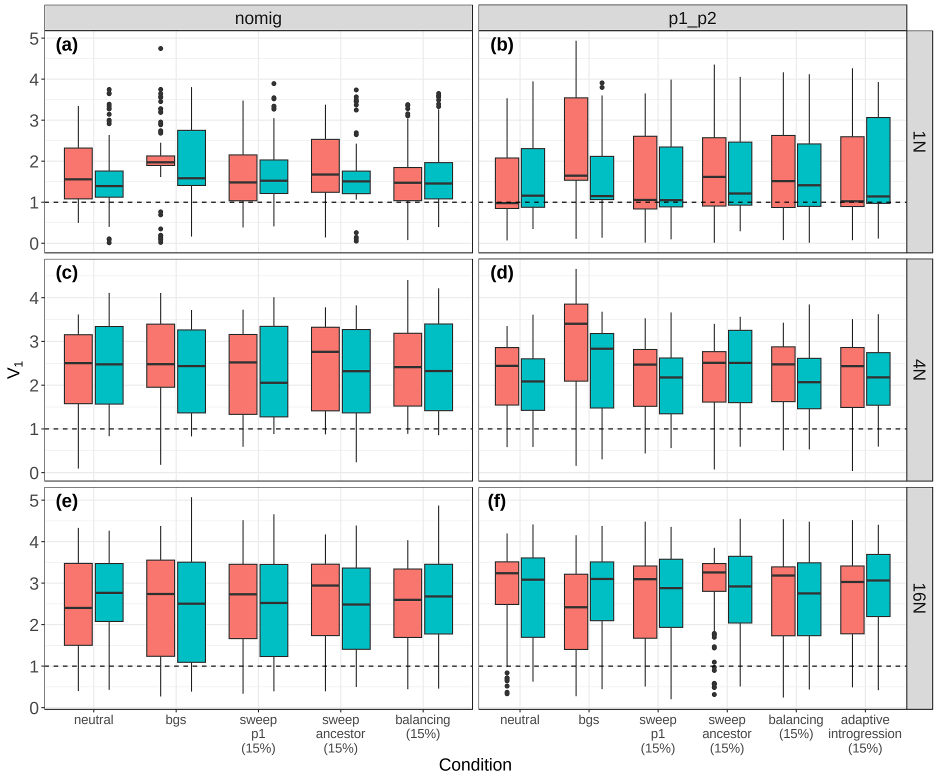


**Supporting Figure S27.** Estimates of $V_1$ in fastsimcoal2. $V_1$ is the size of population 1 relative to the ancestral population. The colors indicate the uniform (pink) and complex (blue) genomic architectures. a) results for T=1*N* and the nomig model; b) results for T=1*N* and the p1_p2 model; c) results for T=4*N* and the nomig model; d) results for T=4*N* and the p1_p2 model; e) results for T=16*N* and the nomig model; f) results for T=16*N* and the p1_p2 model.

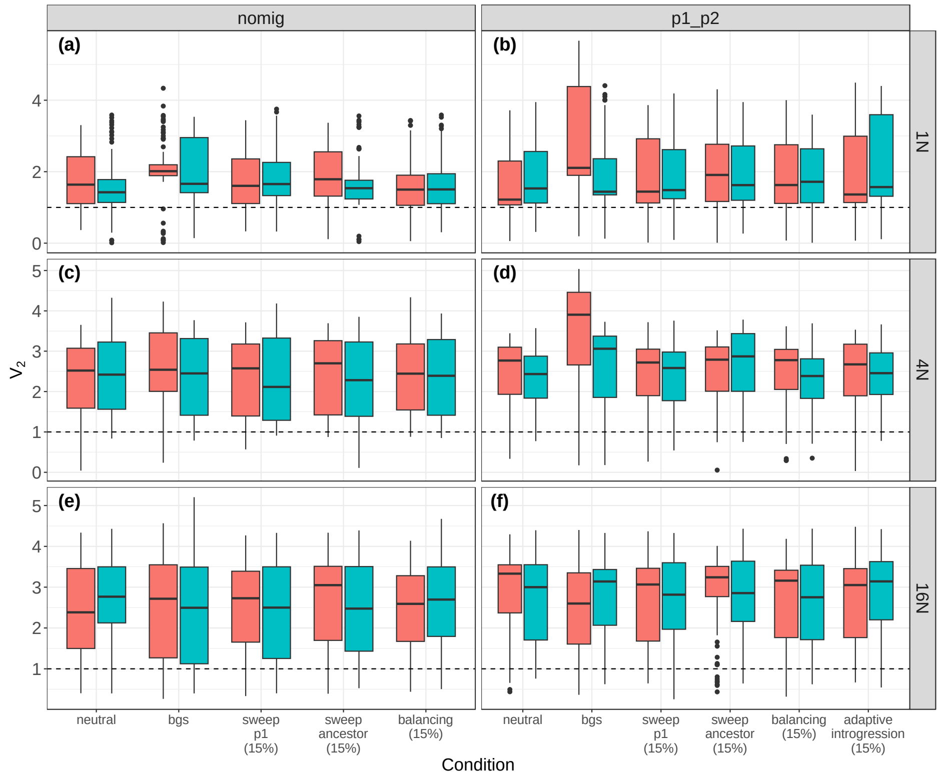


**Supporting Figure S28.** Estimates of $V_2$ in fastsimcoal2. $V_2$ is the size of population 2 relative to the ancestral population. The colors indicate the uniform (pink) and complex (blue) genomic architectures. a) results for T=1*N* and the nomig model; b) results for T=1*N* and the p1_p2 model; c) results for T=4*N* and the nomig model; d) results for T=4*N* and the p1_p2 model; e) results for T=16*N* and the nomig model; f) results for T=16*N* and the p1_p2 model.

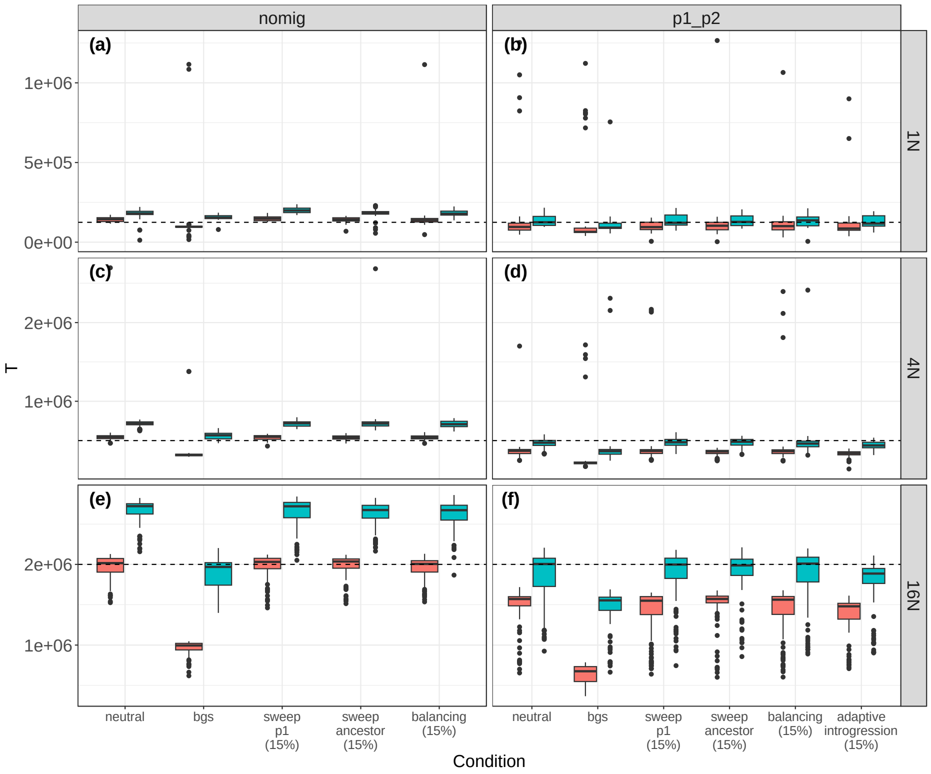


**Supporting Figure S29.** Estimates of *T* in fastsimcoal2. *T* is in units of generations before the present. The colors indicate the uniform (pink) and complex (blue) genomic architectures. a) results for T=1*N* and the nomig model; b) results for T=1*N* and the p1_p2 model; c) results for T=4*N* and the nomig model; d) results for T=4*N* and the p1_p2 model; e) results for T=16*N* and the nomig model; f) results for T=16*N* and the p1_p2 model.

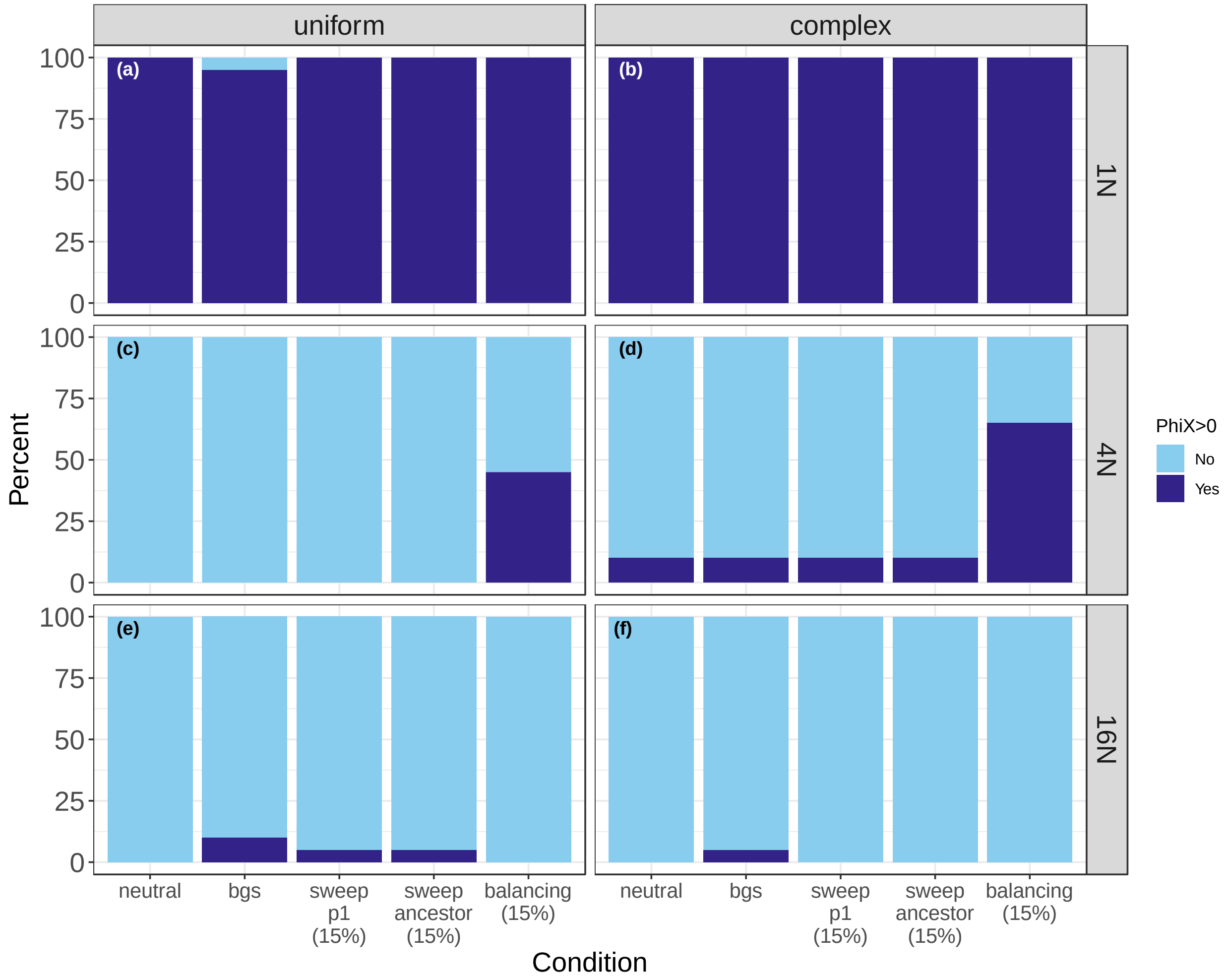


**Supporting Figure S30.** The proportions of replicates with 95% highest posterior density intervals for φ-*X* that do not include zero in the absence of migration in BPP. Light blue indicates cases where the HDI includes zero, and dark blue indicates cases where the HDI does not include zero. a) results for T=1*N* with a uniform genomic architecture; b) results for T=1*N* with a complex genomic architecture; c) results for T=4*N* with a uniform genomic architecture; d) results for T=4*N* with a complex genomic architecture; e) results for T=16*N* with a uniform genomic architecture; f) results for T=16*N* with a complex genomic architecture.

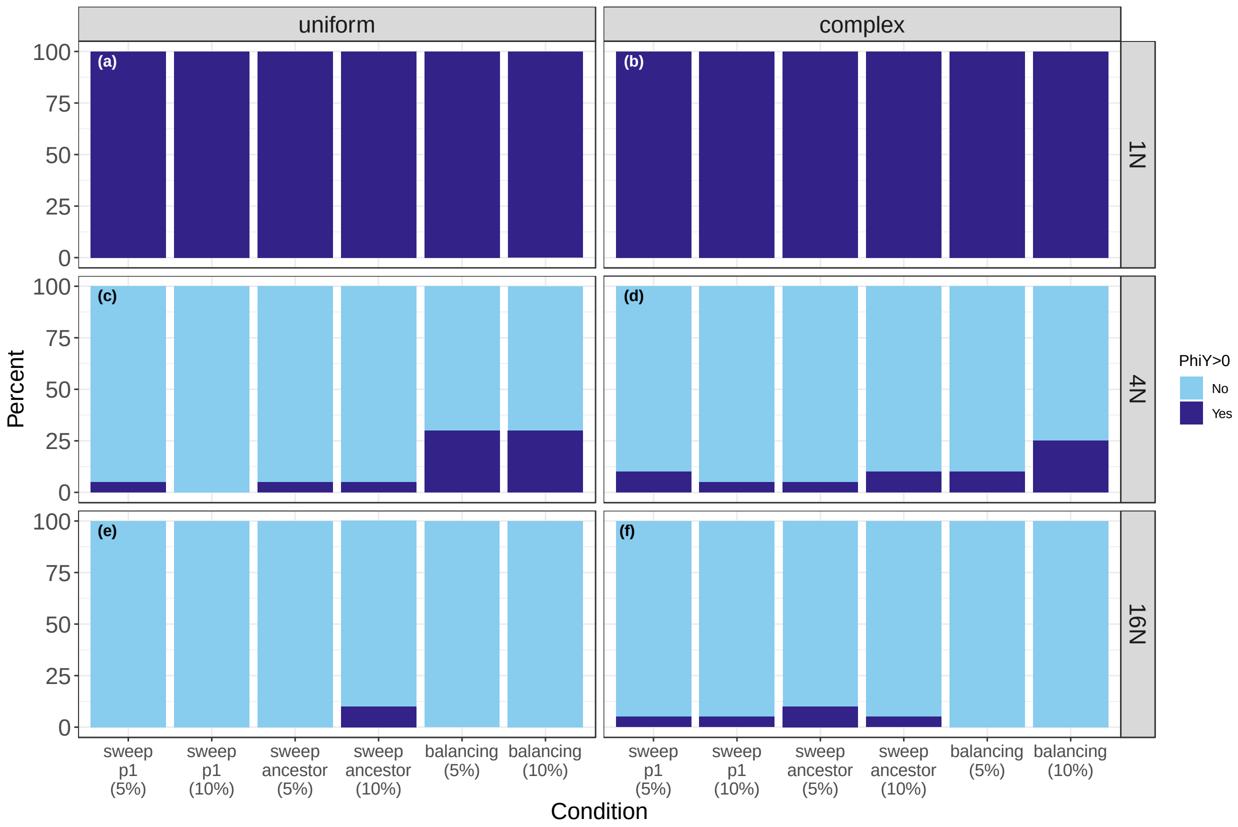


**Supporting Figure S31.** The proportions of replicates with 95% highest posterior density intervals for φ-*Y* that do not include zero in the absence of migration in BPP when sweeps composed 5 or 10 percent of the data. Light blue indicates cases where the HDI includes zero, and dark blue indicates cases where the HDI does not include zero. a) results for T=1*N* with a uniform genomic architecture; b) results for T=1*N* with a complex genomic architecture; c) results for T=4*N* with a uniform genomic architecture; d) results for T=4*N* with a complex genomic architecture; e) results for T=16*N* with a uniform genomic architecture; f) results for T=16*N* with a complex genomic architecture.

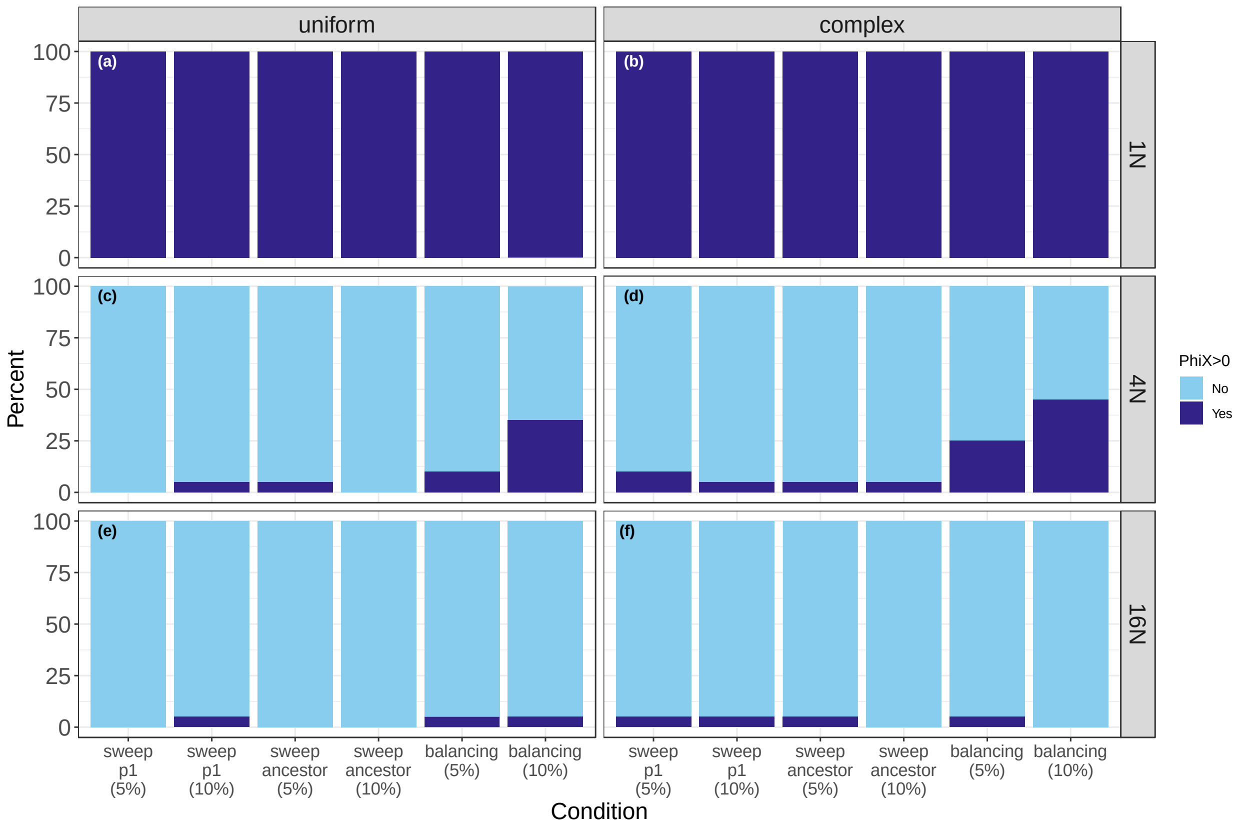


**Supporting Figure S32.** The proportions of replicates with 95% highest posterior density intervals for φ-*X* that do not include zero in the absence of migration in BPP when sweeps composed 5 or 10 percent of the data. Light blue indicates cases where the HDI includes zero, and dark blue indicates cases where the HDI does not include zero. a) results for T=1*N* with a uniform genomic architecture; b) results for T=1*N* with a complex genomic architecture; c) results for T=4*N* with a uniform genomic architecture; d) results for T=4*N* with a complex genomic architecture; e) results for T=16*N* with a uniform genomic architecture; f) results for T=16*N* with a complex genomic architecture.

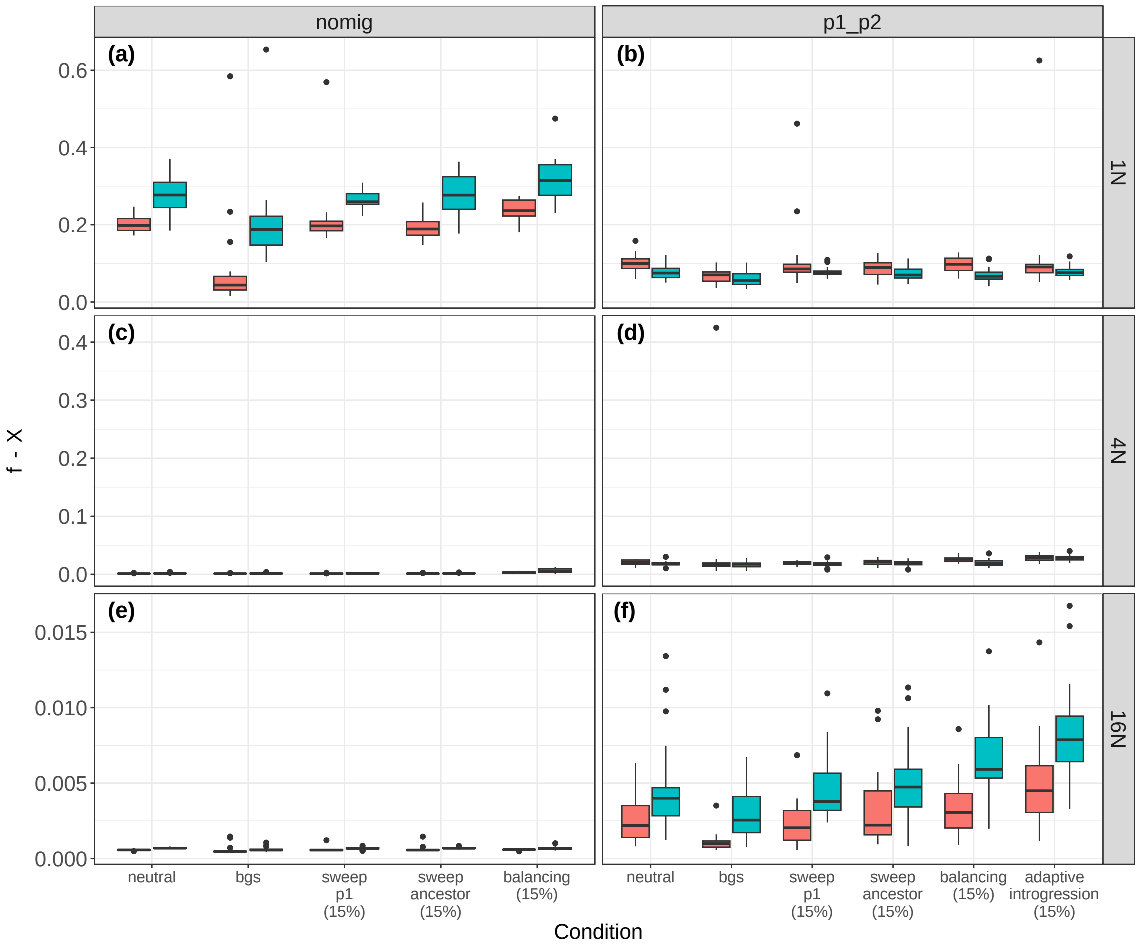


**Supporting Figure S33.** Mean posterior estimates of φ-*X* in BPP. φ-*X* is the weight of the introgression edge X in the MSci model. The colors indicate the uniform (pink) and complex (blue) genomic architectures. a) results for T=1*N* and the nomig model; b) results for T=1*N* and the p1_p2 model; c) results for T=4*N* and the nomig model; d) results for T=4*N* and the p1_p2 model; e) results for T=16*N* and the nomig model; f) results for T=16*N* and the p1_p2 model.

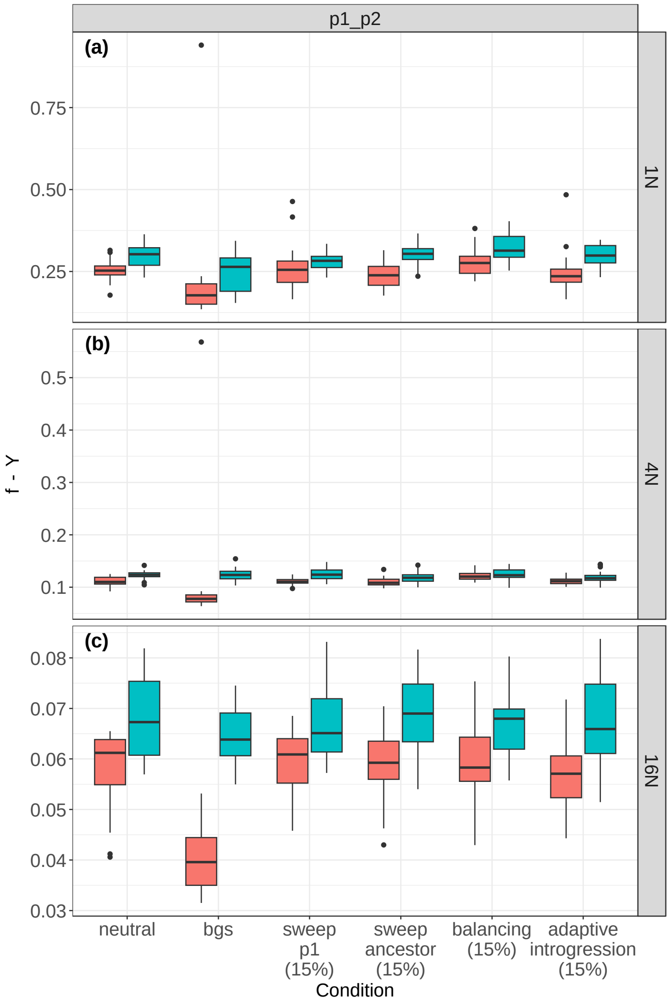


**Supporting Figure S34.** Mean posterior estimates of φ-*Y* in BPP for the migration models. φ-*Y* is the weight of the introgression edge Y in the MSci model. The colors indicate the uniform (pink) and complex (blue) genomic architectures. a) results for T=1*N* and the p1_p2 model; b) results for T=4*N* and the p1_p2 model; c) results for T=16*N* and the p1_p2 model.

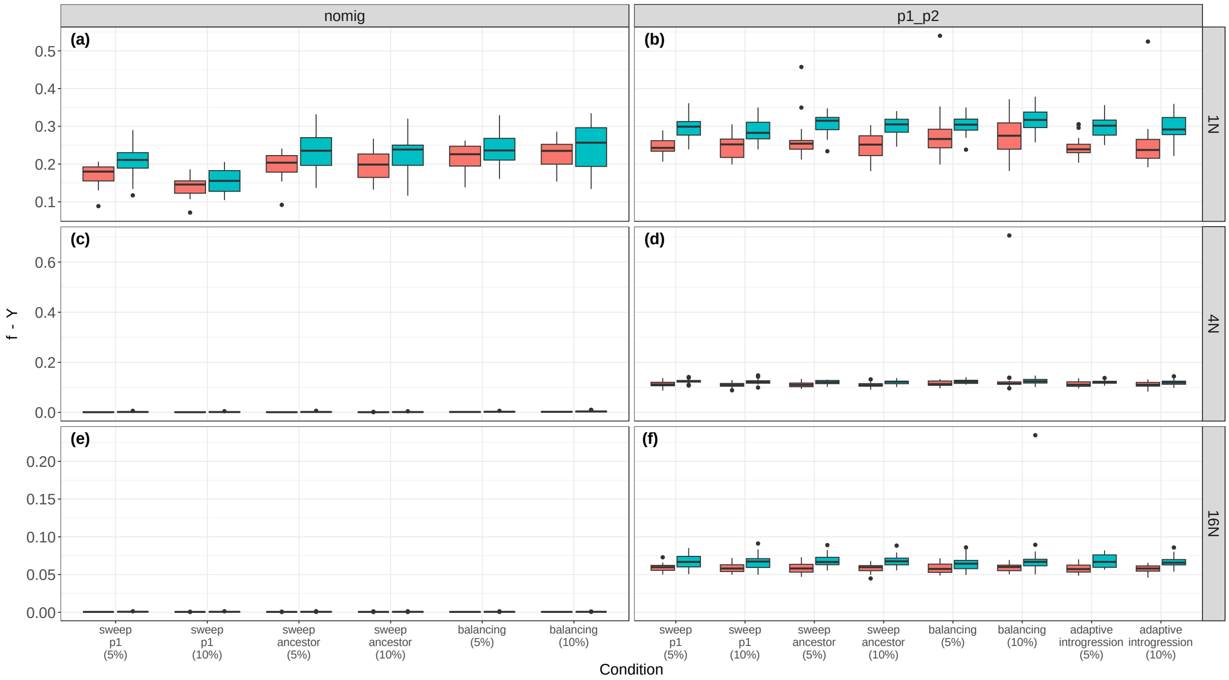


**Supporting Figure S35.** Mean posterior of φ-*Y* in BPP when sweeps composed 5 or 10 percent of the data. φ-*Y* is the weight of the introgression edge Y in the MSci model. The colors indicate the uniform (pink) and complex (blue) genomic architectures. a) results for T=1*N* and the nomig model; b) results for T=1*N* and the p1_p2 model; c) results for T=4*N* and the nomig model; d) results for T=4*N* and the p1_p2 model; e) results for T=16*N* and the nomig model; f) results for T=16*N* and the p1_p2 model.

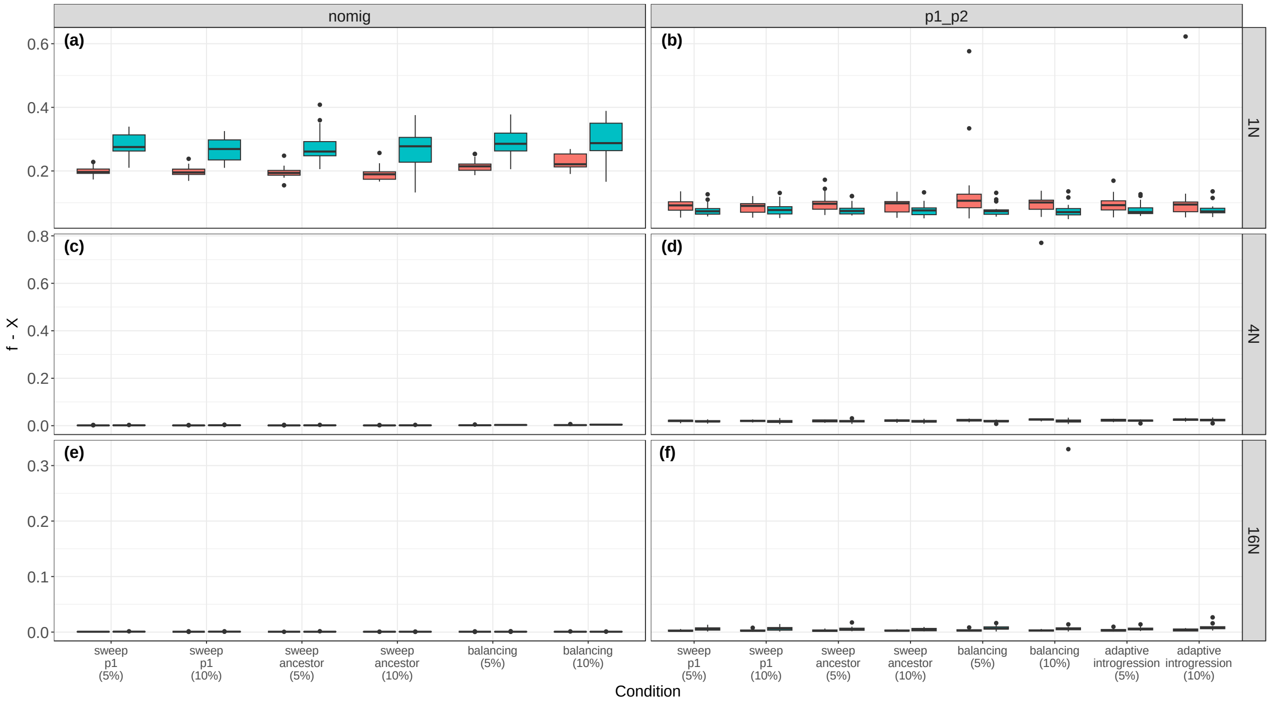


**Supporting Figure S36.** Mean posterior estimates of φ-*X* in BPP when sweeps composed 5 or 10 percent of the data. φ-*X* is the weight of the introgression edge X in the MSci model. The colors indicate the uniform (pink) and complex (blue) genomic architectures. a) results for T=1*N* and the nomig model; b) results for T=1*N* and the p1_p2 model; c) results for T=4*N* and the nomig model; d) results for T=4*N* and the p1_p2 model; e) results for T=16*N* and the nomig model; f) results for T=16*N* and the p1_p2 model.

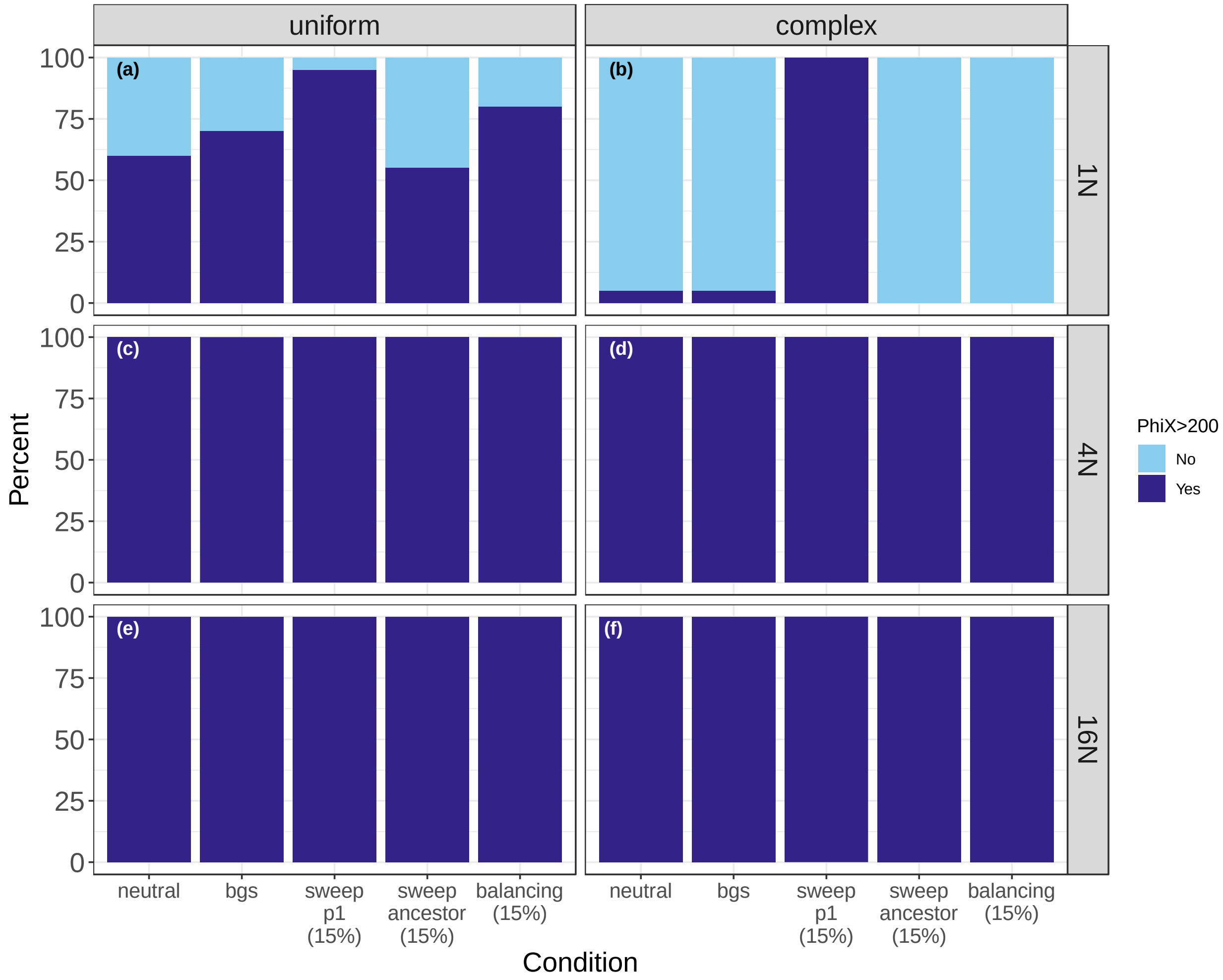


**Supporting Figure S37.** The proportions of replicates with ESS > 200 for φ-X for datasets without migration. Light blue indicates cases where ESS < 200, and dark blue indicates cases where ESS > 200. a) results for T=1*N* with a uniform genomic architecture; b) results for T=1*N* with a complex genomic architecture; c) results for T=4*N* with a uniform genomic architecture; d) results for T=4*N* with a complex genomic architecture; e) results for T=16*N* with a uniform genomic architecture; f) results for T=16*N* with a complex genomic architecture.

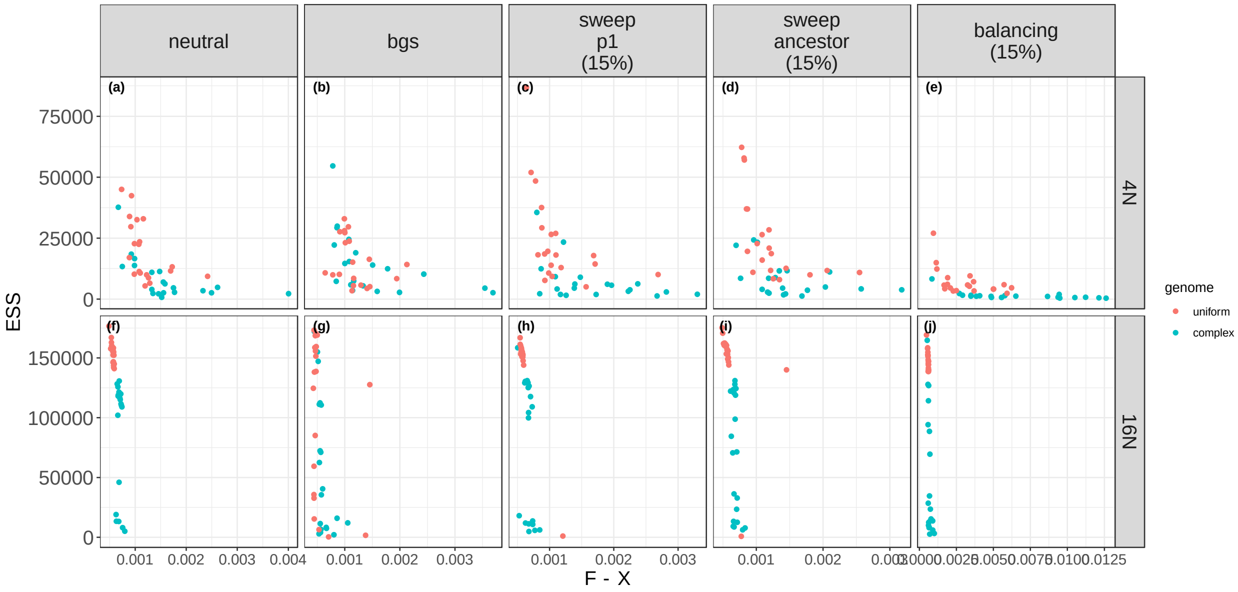


**Supporting Figure S38.** Correlations between the ESS for φ-X and φ-X for datasets without migration. a) results for T=4*N* under the neutral condition; b) results for T=4*N* under the bgs condition; c) results for T=4*N* when 15% of loci experienced a selective sweep in p1; d) results for T=4*N* when 15% of loci experienced a sweep in the ancestral population; e) results for T=4*N* when 15% of loci experienced balancing selection; f) results for T=16*N* under the neutral condition; g) results for T=16*N* under the bgs condition; h) results for T=16*N* when 15% of loci experienced a selective sweep in p1; i) results for T=16*N* when 15% of loci experienced a sweep in the ancestral population. j) results for T=16*N* when 15% of loci experienced balancing selection.

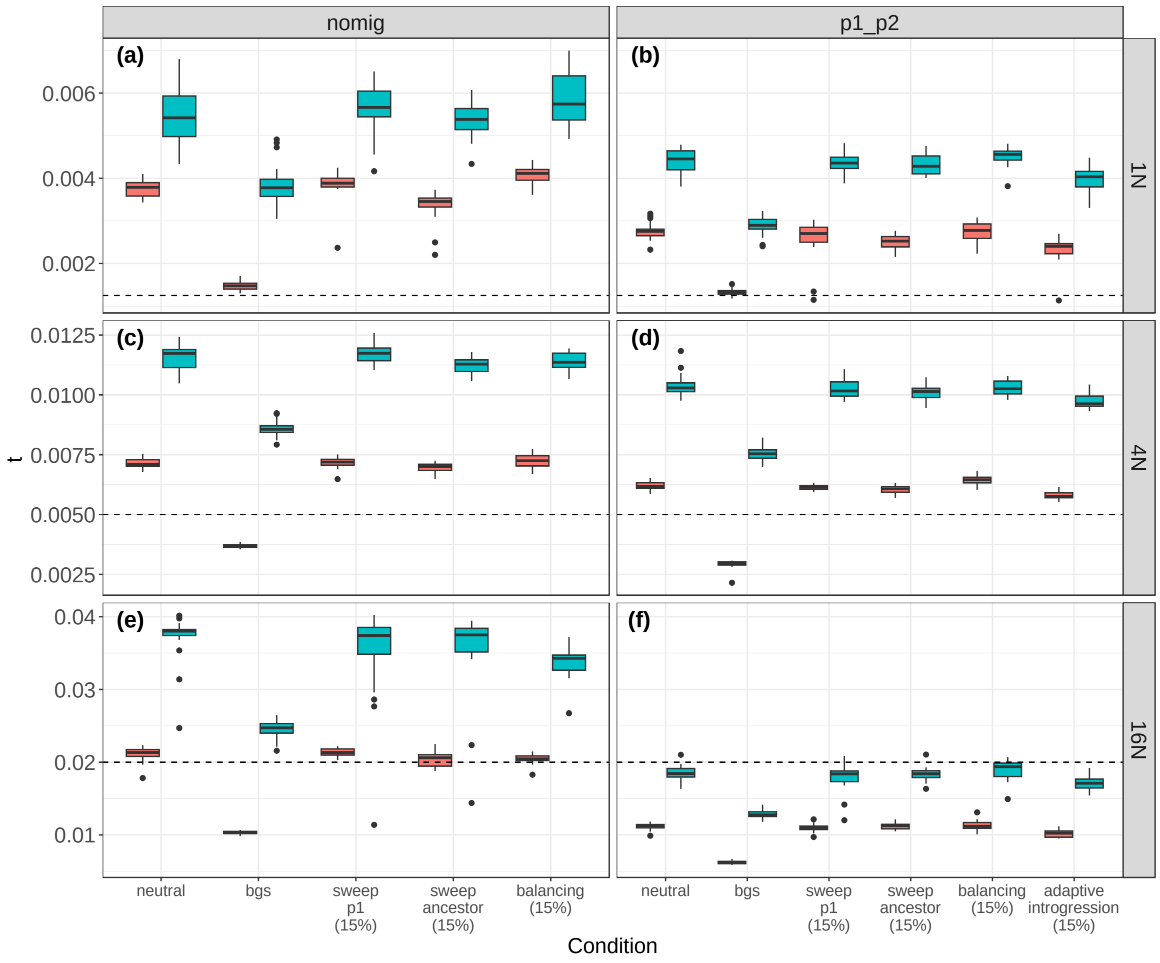


**Supporting Figure S39.** Mean posterior estimates of τ in BPP. τ is in units of expected substitutions per site. The colors indicate the uniform (pink) and complex (blue) genomic architectures. a) results for T=1*N* and the nomig model; b) results for T=1*N* and the p1_p2 model; c) results for T=4*N* and the nomig model; d) results for T=4*N* and the p1_p2 model; e) results for T=16*N* and the nomig model; f) results for T=16*N* and the p1_p2 model.

| Model | Condition | replicate | BF (evidence against the simple model) |
|---|---|---|---|
| **nomig** | bgs | 1 | 293.01 |
| **nomig** | bgs | 2 | -783.87 |
| **nomig** | bgs | 3 | 291.76 |
| **nomig** | bgs | 4 | -76.53 |
| **nomig** | bgs | 5 | -1237.19 |
| **nomig** | neutral | 1 | -726.87 |
| **nomig** | neutral | 2 | 62.34 |
| **nomig** | neutral | 3 | -3.17 |
| **nomig** | neutral | 4 | -484.46 |
| **nomig** | neutral | 5 | 236.28 |
| **p1_p2** | bgs | 1 | 466.37 |
| **p1_p2** | bgs | 2 | 476.97 |
| **p1_p2** | bgs | 3 | -358.27 |
| **p1_p2** | bgs | 4 | 29.91 |
| **p1_p2** | bgs | 5 | 559.58 |

**Supporting Table S1:** Results from Bayes Factor calculations in BPP. Results are from the complex genomic architecture and T=16*N*.
